# Pinpointing of cysteine oxidation sites in vivo by high-resolution proteomics reveals mechanism of redox-dependent inhibition of STING

**DOI:** 10.1101/2020.03.25.008920

**Authors:** Natalia Zamorano Cuervo, Audray Fortin, Elise Caron, Stéfany Chartier, Nathalie Grandvaux

**Affiliations:** CRCHUM - Centre Hospitalier de l’Université de Montréal, 900 rue Saint Denis, Montréal, H2X 0A9, Québec, Canada; Department of Biochemistry and Molecular Medicine, Faculty of Medicine, Université de Montréal, Montréal, H3C 3J7, Québec, Canada

**Keywords:** STING, oxidation, redox, antiviral, inflammation, cancer, autoimmunity, cGAS, proteomics, Cysteine

## Abstract

Protein function is regulated by post-translational modifications, among which reversible oxidation of Cys (Cys ox-PTM) emerged as a key regulatory mechanism of cellular responses. The redox regulation of virus-host interactions is well documented, but in most cases, proteins subjected to Cys ox-PTM remain unknown. The identification of Cys ox-PTM sites in vivo is essential to underpin our understanding of the mechanisms of the redox regulation. In this study, we present a proteome-wide identification of reversible Cys ox-PTM sites in vivo during stimulation by oxidants using a maleimide-based bioswitch method coupled to mass spectrometry. We identified 2720 unique Cys ox-PTM sites encompassing 1473 proteins with distinct abundance, location and functions. Label-free quantification (LFQ)-based analysis revealed the enrichment of ox-PTM in numerous pathways, many relevant to virus-host interaction. Here, we focused on the oxidation of STING, the central adaptor of the innate immune type I interferon pathway induced upon detection of cytosolic DNA. We provide the first in vivo demonstration of reversible oxidation of Cys^148^ and Cys^206^ of STING. Molecular analyses led us to establish a new model in which Cys^148^ oxidation is constitutive, while Cys^206^ oxidation is inducible by oxidative stress or by the natural ligand 2’3’-cGAMP. We show that oxidation of Cys^206^ has an inhibitory function to prevent STING hyperactivation through the constraint of a conformational change associated with the formation of inactive polymers containing intermolecular disulfide bonds. This provides new ground for the design of therapies targeting STING relevant to autoinflammatory disorders, immunotherapies and vaccines.

**Brief summary of the main results:** The function of proteins is regulated by post-translational modifications, among which reversible oxidation of Cys recently emerged as a key component. Comprehension of redox regulation of cellular responses requires identification of specific oxidation sites in vivo. Using a bioswitch method to specifically label Cys subjected to reversible oxidation coupled to mass spectrometry, we identified thousands of novel oxidation sites. Many are relevant to virus-host interaction pathways. Here, we focused on the oxidation of STING, an adaptor critical for activating the innate immune type I interferon pathway engaged upon cytosolic DNA sensing. Molecular studies led us to establish a new model in which STING Cys^148^ is oxidized at basal levels, while Cys^206^ oxidation is induced by oxidative stress and ligand binding. We show that oxidation of Cys^206^ has an inhibitory function to prevent STING hyperactivation. This study provides ground for novel research avenues aimed at designing therapeutics that target this pathway.

## Introduction

Changes of redox status take place during various cellular processes, including virus infections. While chronically elevated levels of reactive oxygen species (ROS) might be associated with oxidative damage, local changes in ROS levels act as redox switches regulating cellular signalling. Increased ROS levels has been documented during various infections by DNA viruses, such as Epstein–Barr virus (EBV) and Kaposi’s sarcoma associated herpesvirus (KSHV), and RNA viruses, including hepatitis C virus (HCV), respiratory syncytial virus (RSV), influenza virus (IAV) and human immunodeficiency virus (HIV) (*1-6*). ROS regulate various aspects of virus-host interactions, including virus replication, host defense and pathogenesis (*5*). Although redox-sensitive virus-induced pathways have started to be identified, such as those leading to Nuclear factor kappa-light-chain-enhancer of activated B cells (NF-κB), Interferon regulatory factor (IRF)-3 or Nuclear factor erythroid 2-related factor 2 (Nrf2) activation, the exact cellular targets of ROS remain poorly characterized (*1, 2, 5*).

ROS act as second messengers in cellular signaling through reversible oxidative post-translational modifications of cysteine residues (Cys ox-PTMs) that allow modulation of protein structure, interaction with protein partners and ligand, localization and activity (*7-9*). Reversible Cys ox-PTMs consist of a variety of modifications, the most widely studied being S-sulfenylation (Cys-SOH), S-glutathionylation (Cys-SSG) and disulfide (S-S) (*7*). Iodoacetamide- or maleimide (Mal)-based bioswitch methods, allowing labeling of oxidized Cys independently of the type of reversible oxidation, coupled to immunoblot or mass spectrometry (MS) have dramatically evolved and made it possible to identify numerous proteins subjected to redox regulation in vivo (*10-14*). However, identification of the corresponding Cys residues subjected to ox-PTM by MS has proven to be challenging and consequently, except in the case of very abundant or highly reactive redox proteins, most often relies on the analysis of the recombinant protein oxidized in vitro (*7, 12, 13, 15, 16, 17, 18-21*).

In our quest to identify novel redox regulated proteins involved in virus-host interaction, our goal was to perform proteome wide identification of in vivo Cys ox-PTM sites upon oxidants stimulation. For this purpose, we used a proven bioswitch method based on the modification of reversibly oxidized Cys with a molecule of Mal-(Polyethylene glycol)2-Biotin (Mal-PEG_2_-Bio) (*11, 22*) followed by an In-solution trypsinization with Peptide Extraction (InsPEx) workflow for identification of labeled peptides by Liquid Chromatography-Tandem Mass Spectrometry (LC-MS/MS). Using this strategy, we successfully identified 2720 unique Cys ox-PTM sites corresponding to 1473 proteins with distinct abundance, localization and functions. Label-free quantification (LFQ)-based analysis pinpointed enrichment of reversible Cys ox-PTMs in numerous functional pathways, including many relevant to virus-host interaction.

Here, we focused on the Cys ox-PTMs of STING, also known as TMEM173, ERIS, MITA and MPYS (*23-26*), the central adaptor and signal relay in the interferon (IFN) pathway dependent on cyclic GMP-AMP synthase (cGAS) (*27-32*). The main function of cGAS-mediated STING activation is innate immune defense when detecting DNA from pathogens, including DNA viruses (*33*). Increasing evidence also support paradoxical roles of the cGAS-STING pathway in tumor immunity and proliferation of tumor cells (*34-36*). STING must be strictly regulated because constitutive activation leads to autoimmune and autoinflammatory disorders (*37-39*). Double strand DNA (dsDNA) released into the cytoplasm are recognized by cGAS leadin g to the production of the 2’3′-cGAMP second messenger that has high-affinity for STING located at the membrane of the endoplasmic reticulum (ER). Binding of 2’3’-cGAMP induces a conformational change and the translocation to a perinuclear compartment (*23, 25, 29, 40*), where it colocalizes and interacts with the TBK1 kinase to ultimately activate the transcription factor IRF3 to induce type I IFNs and other cytokines (*23, 41-43*).

We demonstrate in vivo reversible oxidation of Cys^148^ and Cys^206^ of STING. Cys^148^ oxidation status is independent of cell treatment, while induction of oxidative stress or stimulation with the natural STING ligand 2’3’-cGAMP is required to trigger reversible oxidation of Cys^206^. Further mutational analyses combined to in silico molecular modeling allowed us to establish a novel model in which Cys^206^ oxidation provokes a conformational change associated with formation of inactive diS-containing polymers to prevent STING hyperactivation. This study paves the way to the design of novel STING agonists or antagonists for treatment of autoinflammatory disorders, immunotherapies and vaccines.

## Results

### Proteome-wide identification of Cys ox-PTM sites in basal and oxidant-stimulated U937 cells

Accurate identification of sites of Cys ox-PTM in vivo in mammalian cells in a proteome-wide fashion by Mass spectrometry (MS) has proved to be challenging due to lack of sensitivity (*14, 44*). Here, we used a proven bioswitch method with a Maleimide-(Polyethylene Glycol)2-Biotin (Mal-PEG_2_-Bio) probe to specifically label Cys subjected to reversible ox-PTM independently of the oxidation type (**Fig. 1A**) (*11, 17, 45*). In pilot experiments, Mal-PEG_2_-Bio-labeled proteins generated from cells stimulated for 20min with 0.5mM diamide, a thiol-oxidizing agent (*46*), were subjected to a standard procedure of NeutrAvidin-mediated pull-down (*47*), followed by in-gel trypsinization before analysis by LC-MS/MS (In-gel trypsinization workflow, **fig. S1A-B**). The fragmentation spectra of the detected peptides were searched on Mascot and Scaffold for peptides containing Cys with a mass shift of 525.23 Da corresponding to the molecular weight (MW) of the Mal-PEG_2_-Bio moiety (**Fig. 1C**). While this strategy permitted identification of proteins subjected to Cys ox-PTM sites, the ratio of peptides containing at least one biotinylated Cys against the total number of peptides was as low as 0.5% (**fig. S1C**). Moreover, assignment of peptides to human proteins using the same programs revealed that they corresponded only to well-known redox proteins, such as thioredoxins (Trxs), peroxiredoxins (Prxs) and glutathione peroxidases (GPxs) (*48, 49*) (**Data File S1**). As an attempt to reach a better sensitivity in the detection of Cys ox-PTM sites, a modified workflow was used. Mal-PEG_2_-Bio-labelled proteins were immediately subjected to in-solution trypsin digestion, followed by a solid-phase extraction step using reverse-phase tC18 SepPak liquid-solid-phase cartridge before NeutrAvidin pull-down and elution in 8M guanidine/HCl pH 1.5 (In-solution trypsinization with Peptide Extraction (InsPEx) workflow, **Fig. 1B**). Using this InsPEx workflow, Mal-PEG_2_-Bio-modified peptides were found highly enriched to a ratio over total peptides of 83% (**fig. S1C and Data File S2**). Importantly, the identified peptides containing at least one biotinylated Cys encompassed proteins with a diverse range of activities and localization (**Data File S2**).

**Fig. 1.**
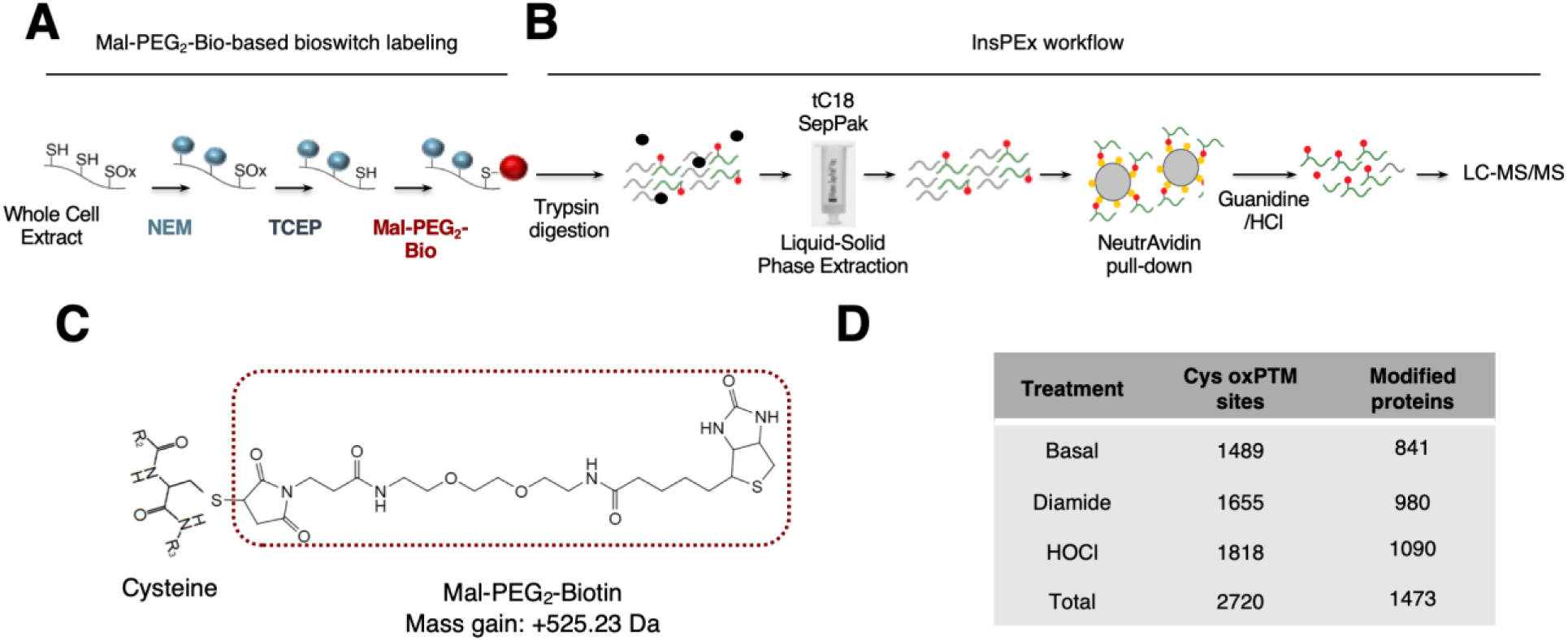
Identification of reversible Cys ox-PTMs induced by oxidants in vivo using unbiased Mal-PEG_2_-Bio-based bioswitch labeling strategy coupled to mass spectrometry. **A-B**. Schematic of the workflow used to label reversible Cys ox-PTMs with the Mal-PEG_2_-Bio bioswitch method (**A**) followed by In-solution trypsinization with peptide extraction (InsPEx, **B**) used to generate and isolate peptides containing at least one Mal-PEG_2_-Bio-labelled Cys for identification by LC-MS/MS. In (**A**), Cys-SH residues were specifically alkylated with N-ethylmaleimide (NEM). Reversibly oxidized Cys (SOx) were then reduced using tris(2-carboxyethyl)phosphine (TCEP) before labeling with EZ-Link Maleimide-PEG_2_-Biotin (Mal-PEG_2_-Bio). In (**B**), proteins from (**A**) were digested in solution using trypsin and peptides were subjected to liquid-solid phase extraction followed by a NeutrAvidin pull down and analysis by LC-MS/MS. **C**. Schematic of a modified Cys residue carrying a Mal-PEG_2_-Bio moiety that adds a mass of 525.23 Da to the Cys. **D**. U937 cells were left untreated or stimulated with 0.5mM diamide or 100µM hypochlorous acid (HOCl) for 20min and 10min, respectively. Whole Cell Extracts were subjected to the workflow described in **A-B**. Table showing total number of Mal-PEG_2_-Bio-labelled Cys sites and the number of corresponding proteins identified in each condition following the InsPEx method. Results of three independent experiments were combined for identification of the Mal-PEG_2_-Bio-labeled peptides.

Based on these pilot experiments, the InsPEx workflow was pursued to identify novel proteins involved in virus-host interaction susceptible to reversible Cys oxidation in vivo. U937 cells unstimulated (basal conditions) or treated with 0.5mM diamide or 100µM hypochlorous acid (HOCl), an oxidant produced in vivo during infections as a result of neutrophils activity (*50*), for 20min and 10min respectively, were subjected to Mal-PEG_2_-Bio-based bioswitch labeling followed by the InsPEx workflow. Peptides extracted from triplicate independent experiments were analyzed by LC-MS/MS. Results of triplicates were combined for identification of the Mal-PEG_2_-Bio-labeled peptides. Overall, this strategy led to the identification of 2699 unique peptides containing 2720 biotinylated Cys residues. Amongst these, 1489 Cys ox-PTM sites belonging to 841 different proteins were found at basal levels; 1655 Cys ox-PTMs corresponding to 980 proteins were identified in cells stimulated with diamide; 1818Cys ox-PTMs encompassing 1090 proteins were found in HOCl-stimulated cells (**Fig. 1D and Data File S3**).

### Identification of pathways related to virus-host interaction enriched in proteins subjected to Cys ox-PTMs

To get insight into the functional implication of the proteins found to contain Cys ox-PTMs under basal or oxidative stress conditions, a Label Free Quantification (LFQ) method was applied considering only Mal-PEG_2_-Bio-labeled Cys identified in at least two out of the three independent replicates per condition to eliminate possible “false positive” hits (*51, 52*). The accuracy of our approach was first validated by the identification of Cys ox-PTM sites previously documented to be oxidized in vitro or in vivo (**fig. S2**).

Next, Gene Ontology (GO) enrichment analysis was performed using the BiNGO plugin into Cytoscape (*53, 54*). To look for Cys ox-PTMs occurring exclusively at basal levels or in oxidative stress condition independently of the type of oxidant, diamide and HOCl conditions were pooled in the analysis. A total of 74 biological processes (BP) were enriched in basal state and 212 in oxidant-stimulated cells (see **Data File S4** for a complete list). Several enriched BPs were relevant to host-virus interactions (**Fig. 2A**): in unstimulated cells, they include translation related processes (translation elongation and translation), and several processes associated with the host immune defense (immune effector process, immunoglobulin biosynthetic process, leukocyte mediated cytotoxicity, immunesystem process); upon oxidant stimulation, translation and immune related processes (leukocyte activation, immune system process) were also enriched, in addition to BPs related to intracellular protein localization (intracellular transport, establishment of protein localization, protein transport and nuclear transport). Further identification of functional clusters amongst these enriched BPs was performed through network community analyses using the fast greedy GLay algorithm (**fig. S3A**). This revealed the prevalence of functional clusters related to immune functions (mediated cytotoxic immunity and antigen symbiont stress) at basal levels and to cell transport, translation frameshifting termination, regulation of phosphorylation and immunity in cells stimulated with oxidants. Fourteen molecular functions (MF) were found significantly enriched in basal conditions and 54 upon oxidants stimulation (see **Data File S4** form a complete list). One of the most enriched MF, both at basal levels and under conditions of oxidative stress, is related to the catalytic functions of proteins (**Fig. 2B**). Further analysis using the GLay algorithm revealed association of these functions into 4 clusters, i.e. peptidase isomerase activity, binding of NAD, structural molecule activity and immune system (MHC class protein), at basal levels (**fig. S3B**). Upon oxidative stress, functional clusters associated with catalytic activities were related to kinase and phosphatase activities (Ser/Thr activity, protein phosphotyrosine binding and adenyl nucleotide ribonucleotide) (**fig. S3B**).

**Fig. 2.**
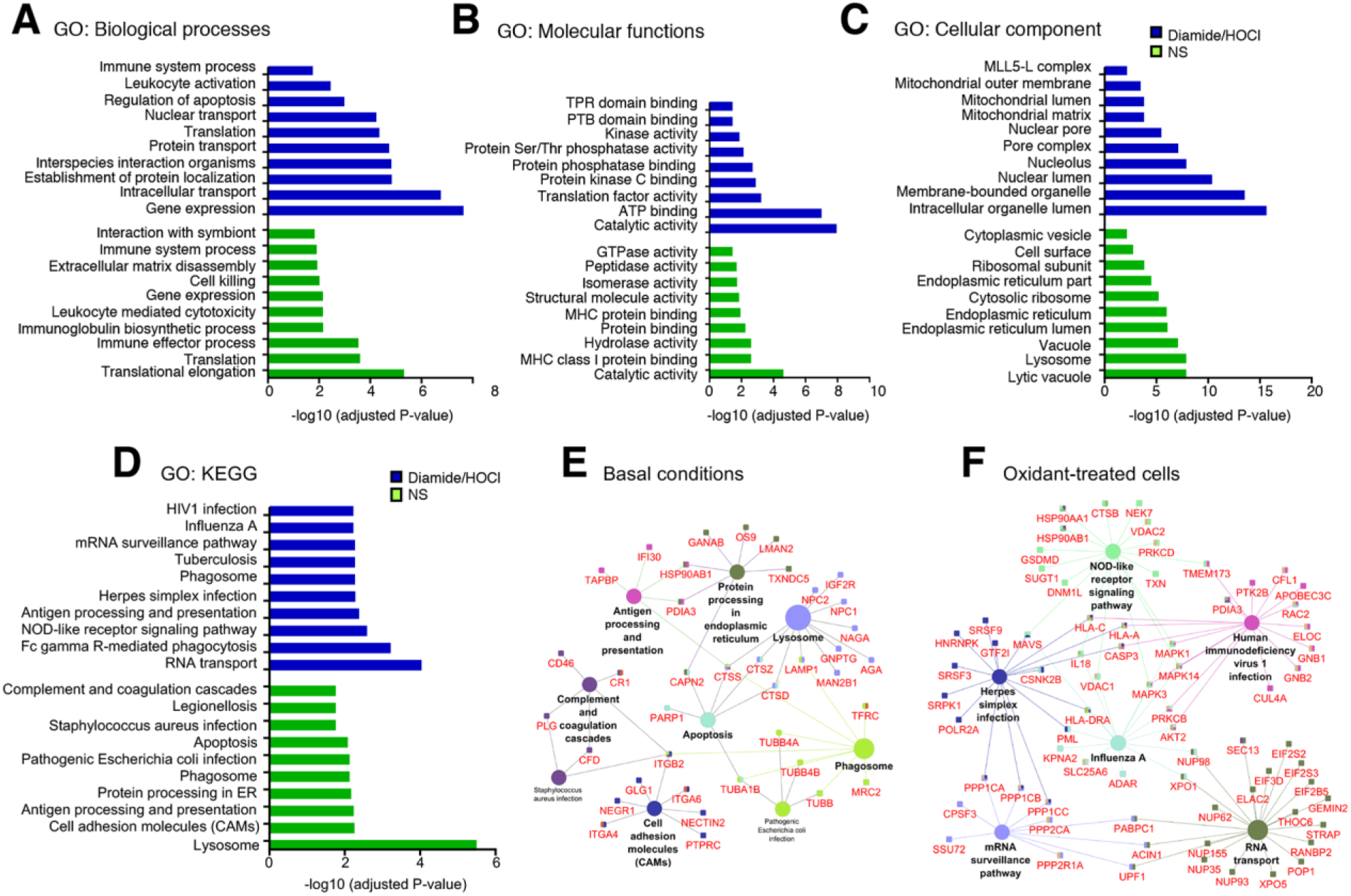
Functional analysis of reversible Cys ox-PTMs induced by oxidants. **A-C**. Gene Ontology (GO) enrichment analysis of identified reversible Cys ox-PTMs in non-stimulated (NS) and oxidant (diamide or HOCl)-stimulated conditions described in **Fig. 1D** based on the biological processes (**A**), molecular functions (**B**) and cellular component (**C**). Selected host-virus interaction relevant terms are shown, and the full list of enriched terms is available in Data File S4. **D-F**. KEGG pathways enrichment analysis of reversible Cys ox-PTMs identified in non-stimulated (basal) and oxidant (diamide/HOCl)-treated conditions described in **Fig. 1D**. Selected host-virus interaction relevant terms are shown in (**D**) and the full list of overrepresented pathways is available in Data File S5. Oxidized proteins of selected KEGG pathways enriched at basal levels (**E**) and in oxidant-treated cells (**F**) are mapped. Only Mal-PEG2-Bio-labeled Cys identified in at least two out of three independent replicates per condition were taken into account fo r the functional analysis.

Analysis of the cellular component (CC) identified 30 and 74 GO terms enriched under basal and oxidants stimulation conditions, respectively (see **Data File S4** for full list). In both conditions, the most enriched compartment corresponds to the cytoplasm (**Data File S4**). At basal levels, terms associated with vesicles/vacuoles (lytic vacuole, lysosome, vacuole and cytoplasmic vesicle), grouped into the functional cluster membrane-bounded vesicle vacuole were highly enriched, as were ER-related terms (ER, ER lumen and ER part) (**Fig. 2C and fig. S3C**). In contrast, following oxidant treatment, the enriched CC terms were corresponded to clusters related to the nucleus (nuclear periphery spliceosome, nucleoplasmic shuttling complex) and mitochondria (mitochondrial part nucleoid) (**Fig. 2C and fig. S3C**).

To further characterize the signaling pathways related to host-virus interaction subjected to redox regulation, Kyoto Encyclopedia of Genes and Genomes (KEGG) pathway analyses were performed using the ClueGO plugin (*55*). At basal levels, 16 KEGG terms were significantly enriched compared to 26 in cells subjected to oxidant treatments (see **Data File S5** for full list and **fig. S4** for full map). Amongst these enriched pathways, several are relevant to virus-host interaction (**Fig. 2D-F**): in unstimulated cells, pathways related to the host defense system, such as those associated with lysosomes, phagosomes, antigen processing and presentation and complement and coagulation cascades (**Fig. 2D and E**); in oxidant-treated cells, pathways related to the immune system, including phagosome, Fc gamma R-mediated phagocytosis, antigen processing and presentation, NOD-like receptor signaling pathway, RNA-associated pathways (RNA transport, mRNA surveillance pathway) and specific virus infections-related pathways (HIV1 infection, Influenza A, Herpes simplex infection) (**Fig. 2D and F**). Together, our data revealed that in a cellular context, hundreds of proteins involved in virus-host interaction undergo reversible Cys ox-PTM, making it possible to envisage new regulatory mechanisms.

### STING Cys^206^ undergoes oxidized reversible ox-PTM upon oxidative stress and 2′3′-cGAMP stimulation

Amongst the Cys ox-PTM-containing proteins related to host-virus interaction, identified in the KEGG pathways enriched in cells subjected to oxidant, is the DNA virus antiviral signaling adaptor STING encoded by TMEM173 (**Fig. 2F**). Oxidation of Cys^206^ was detected in HOCl-stimulated cells (**Fig. 3B and Datafile S3**). Further analysis of our data set also revealed oxidation of Cys^148^, but the modification was detected in both the basal condition and following oxidant treatment (**Fig. 3A-B and Datafile S3**).

**Fig. 3.**
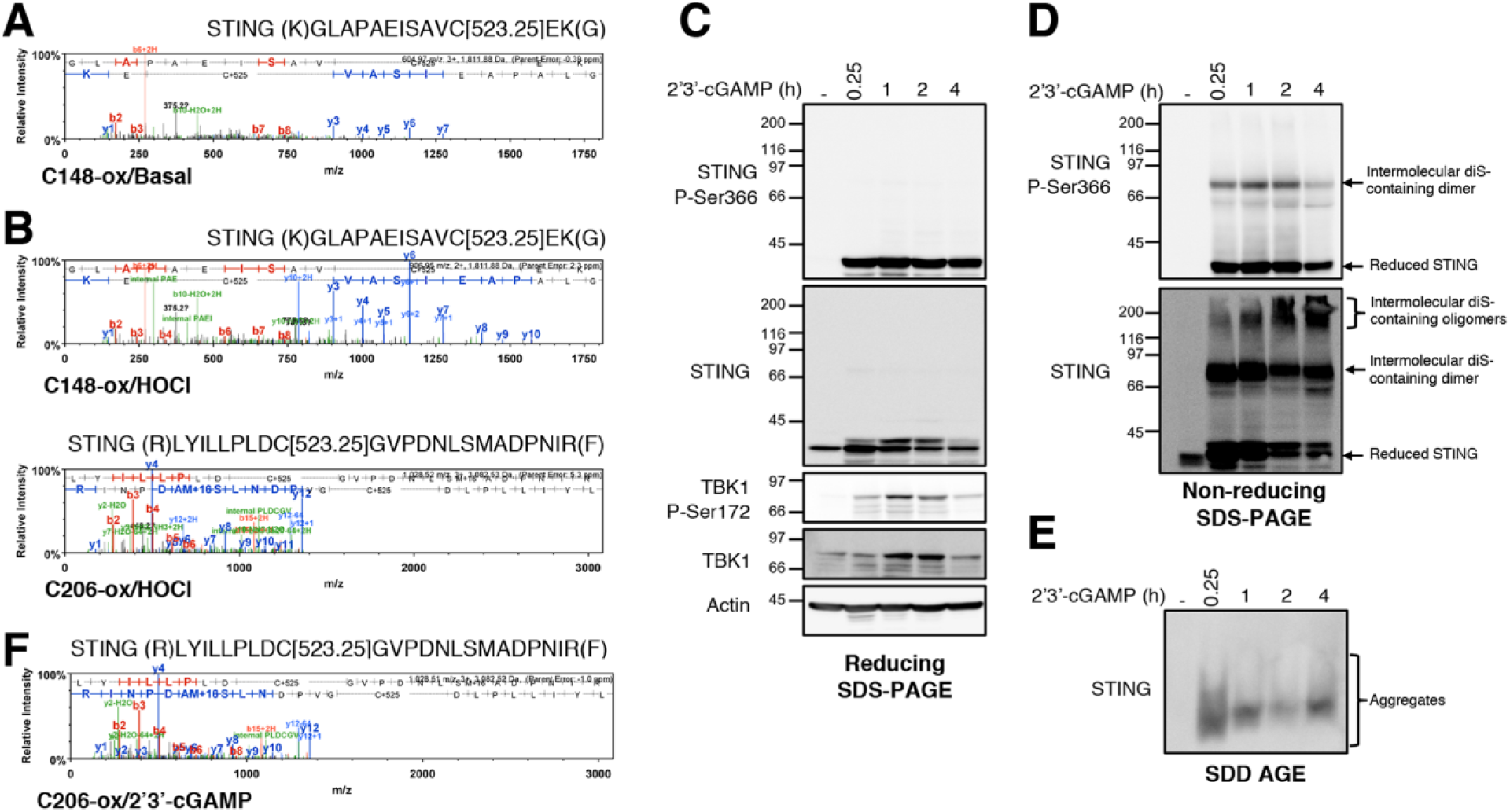
Characterization of STING Cys reversible oxidation upon HOCl and 2’3’-cGAMP stimulation. **A-B**. Fragment spectra of the STING peptides identified in the experiments described in **Fig. 1D** showing the modification of the Cys residues, C148 and C^206^, with Mal-PEG_2_-Bio (+525.23). The fragment picks of both b and y ions are shown in the spectra in red and blue, respectively. **C**. U937 cells were infused with 2’3’-cGAMP at 24µg/mL for the indicated times. Cells were harvested and whole cell extracts were immediately resolved by reducing (with DTT) and non-reducing (without DTT) SDS-PAGE and by SDD-AGE. Proteins were detected by immunoblot using anti-actin, anti-TBK1, anti-phospho-TBK1-Ser^172^, anti-STING and anti-phospho-STING-Ser^366^ antibodies. Molecular weight (kDa) markers are indicated on the left side of the SDS-PAGE gels. The data are representative of three independent experiments. **D**. U937 cells were infused with or without 2’3’-cGAMP at 24µg/mL for 30min. WCE were subjected to the workflow described in **Fig. 1A-B** to identify reversible Cys ox-PTMs. Fragment spectrum of the STING peptide containing the Cys residue C206 modified with Mal-PEG_2_-Bio identified in the 2’3’-cGAMP stimulated condition. All MS/MS spectra were visualized with scaffold, version 4.8.4.

STING is activated through the docking of 2’3’-cGAMP generated upon DNA recognition by cGAS (*27, 29, 31, 32*). Therefore, we next sought to determine whether Cys^206^ also undergoes reversible oxidation following stimulation by 2’3’-cGAMP. U937 cells were first treated with 2’3’-cGAMP for various times and analyzed by immunoblot. Phosphorylation of STING on Ser^366^, a marker of STING activation (*56*), and of the downstream effector kinase TBK1 on Ser^172^ (*57*), were detected after 15min of 2’3’-cGAMP stimulation (**Fig. 3C**). Previous studies have reported that stimulation by 2’3’-cGAMP leads to disulfide (diS) bond-dependent form of STING dimer and high molecular weight (HMW) aggregates (*58, 59*). Analysis of whole cell extracts (WCE) by non-reducing SDS-PAGE confirmed the formation of intermolecular diS-containing dimeric form of STING, migrating at about 74kDa after 15min of stimulation (**Fig. 3D**). It is noteworthy, that additional intermolecular diS-containing oligomeric species of higher MW were also detected during the course of 2’3’-cGAMP stimulation (**Fig. 3D**). Ser^366^ phosphorylation of STING was detected in the diS-containing dimeric form (**Fig. 3D**). Resolution of WCE by SDD-AGE also confirmed the formation of HMW aggregates starting at 15min of 2’3’-cGAMP stimulation (**Fig. 3E**). Based on these observations, U937 cells subjected to stimulation with 2’3’-cGAMP for 30min and the corresponding control cells were subjected to Mal-PEG_2_-Bio-based bioswitch labeling followed by the InsPEx workflow to detect reversible Cys ox-PTMs (**Fig. 1A**). Oxidation of Cys^206^ was detected in 2’3’-cGAMP-stimulated cells, but not in control cells (**Fig. 3F and fig. S5**). Overall, our data unveils the reversible oxidation of Cys^148^ and Cys^206^ of STING. While Cys^148^ oxidation status is independent of cell treatment, oxidative stress induced by HOCl and stimulation with the natural STING ligand 2’3’-cGAMP trigger reversible oxidation of Cys^206^.

### Cys^206^ oxidation dampens STING activation

The importance of Cys^148^ oxidation in the capacity of STING to be activated by 2’3’-cGAMP was recently documented (*60, 61*). Hence, in this study we focused on elucidating the role of Cys^206^ oxidation. According to the available tridimensional structures of human STING dimer bound to 2’3’-cGAMP (*29, 61-65*), Cys^206^ lies in the 2’3’-cGAMP binding pocket (**Fig. 4A-C**). To start investigating the impact of Cys^206^ oxidation, we used the PyTMs plugin in PyMOL software to perform in silico molecular assessment of the steric van-der-Waals (vdW) hindrance in the structure due to Cys^206^ modification by sulfenylation (SOH). Predicted steric structural strains (vdW clashes, (*66*)) were observed within the dimer interface and between Cys^206^ and Arg^284^ (**Fig. 4B and C**) indicating that a change in conformation must take place after oxidative modification of Cys^206^.

**Fig. 4.**
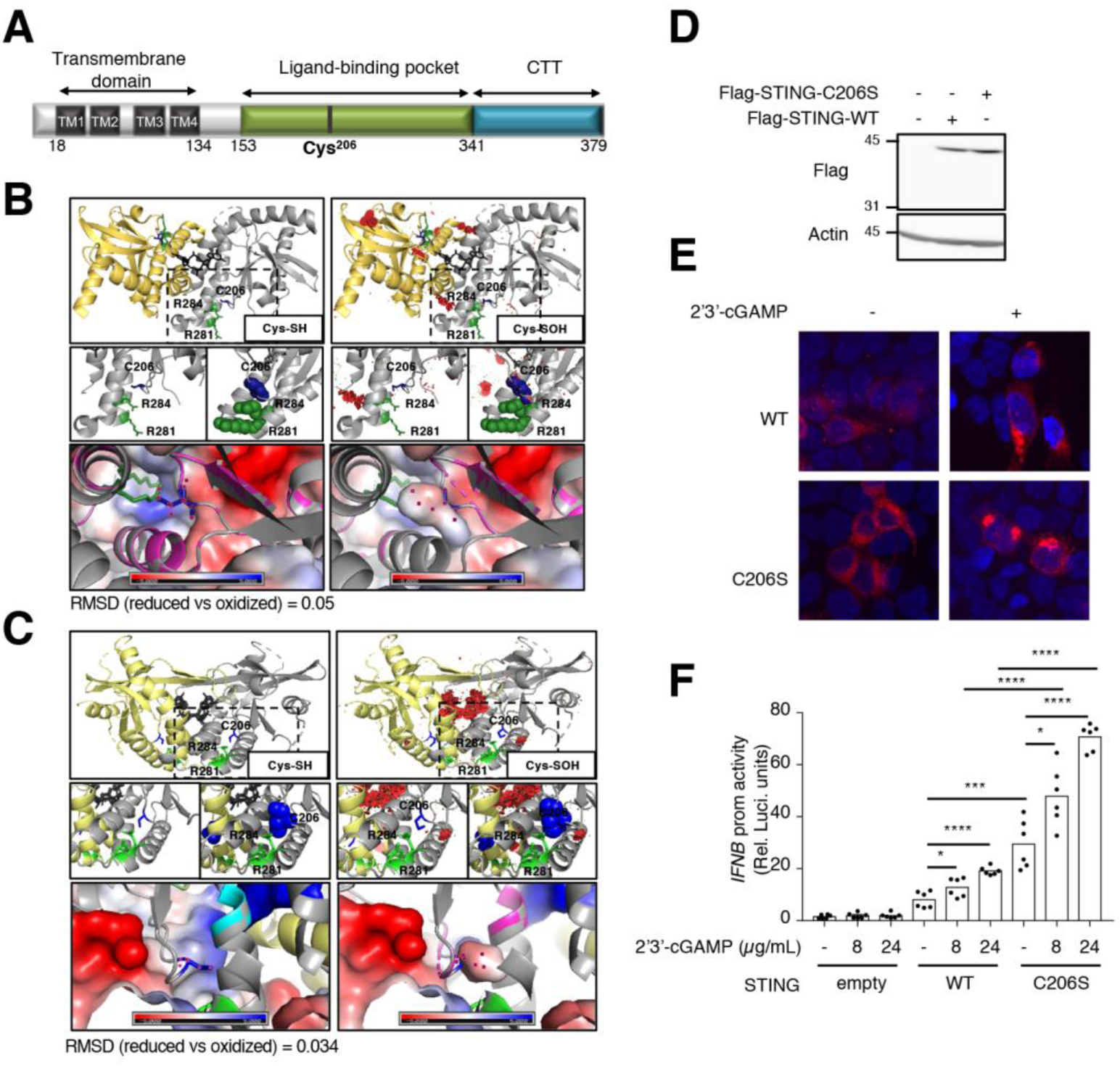
Role of Cys^206^ oxidation on 2’3-cGAMP-activated STING structure and function. **A**. Schematic representation of the linear structure of STING showing the transmembrane domains (TM), the ligand binding pocket and the C terminal tail (CTT). The position of Cys^206^ is indicated. **B-C**. Annotated structures, pdb4ef4 (**B**) (*64*) and pdb6dnk (*61*) **(C)**, of STING bound to a molecule of cyclic di-GMP (in black) are shown. Top panels: structures with reduced Cys^206^ (Cys-SH) are shown on the left side; structures with sulfenylated Cys^206^ (Cys-SOH) modelized using the PyTMs plugin in PyMol are shown on the right side. Middle panels: the zone where the three amino acids are located is zoomed out. Lower panels: APBS Electrostatics plugin was used to visualize the electrostatic surface and superpose the structures containing reduced and oxidized Cys^206^. Cys^206^ (C206) is shown in blue. Arg^281^ (R281) and Arg^284^ (R284) are shown in green. Lateral chains of the three amino acids are represented in sticks. Steric van-der-Waals (vdW) hindrance (vdW clashes) are shown in red. RMSD: root-mean-square deviation of atomic positions. **D-E**. HEK293T cells were transfected with Flag-tagged wild-type (WT) or C206S STING-encoding plasmids or the corresponding control plasmid (empty) before stimulation with 2’3’-cGAMP as indicated. In **D**, whole cell extracts were resolved by standard SDS-PAGE. Proteins were detected by immunoblot using anti-actin and anti-Flag antibodies. Molecular weight (kDa) markers are indicated on the left side of the SDS-PAGE gels. The data are representative of three independent experiments. In **E**, Confocal images of cells immunostained for Flag-STING (anti-DYKDDDDK) and nuclei using 4’ 6, diamido 2 phenylindole (DAPI). Representative of two independent experiments **F**. HEK293T cells transfected with the IFNβ promoter luciferase reporter construct together with the Flag-tagged WT or C206S STING-encoding or the empty plasmid were further transfected with or without 2’3’-cGAMP at the indicated concentration. Relative luciferase activities were measured at 24h post-stimulation. Mean +/- SEM, n=6, unpaired t-test.

To further delineate the role of Cys^206^ reversible ox-PTM in the regulation of STING function, we tested the impact of Cys^206^ mutation into Ser (STING-C206S) (**Fig. 4D**), which suppresses its capacity to be oxidized, on STING-mediated signaling. First, confocal indirect immunofluorescence analysis revealed that Flag-tagged-STING WT and -STING-C206S exhibit similar localization at basal levels and translocation to perinuclear punctate structures upon 2’3’-cGAMP stimulation (**Fig. 4D-E**). Next, the activity of STING WT and STING-C206S was measured through their capacity to transactivate the IFNβ promoter using a luciferase-based reporter assay at basal levels and upon stimulation with 2’3’ cGAMP. As expected, the activity of STING WT was significantly enhanced by 2’3’-cGAMP in a dose-dependent manner (**Fig. 4F**). The STING-C206S mutant strongly induced the IFNβ promoter in the absence of ligand compared to STING WT (29.5 fold compared to 8.1 fold, **Fig. 4F**). STING-C206S mutant activity was highly increased by 2’3’-cGAMP compared to STING WT (70.7 fold compared to 19.2 fold, **Fig 4F**). Altogether, our data suggests that modification of Cys^206^ plays an inhibitory function and support a model in which Cys^206^ oxidation would prevent STING hyperactivation.

Next, we studied the impact of HOCl on STING in the absence or presence of stimulation by 2’3’-cGAMP. For this purpose, cells expressing Flag-tagged-STING WT or - STING-C206S were stimulated or not with 2’3’-cGAMP for 1h before further treatment with HOCl. First, the capacity of STING WT and STING-C206S to activate the IFNβ promoter luciferase reporter was evaluated. Our results indicate that HOCl inhibits 2’3’-cGAMP-induced activity of STING WT by 44%, while inhibition of C206S activity was limited to 22% (**Fig. 5A**). In the absence of 2’3’-cGAMP stimulation, HOCl diminished STING WT activity by 40%, whereas no significant inhibition of STING-C206S mutant was observed (**Fig. 5B**). These observations confirmed that oxidative stress inhibits STING activation in a Cys^206^-dependent manner. In order to gain further insight into the mechanism by which Cys206 oxidation inhibits STING activity, phosphorylation of STING on Ser^366^ was analyzed by immunoblot. Stimulation by 2’3’-cGAMP led to hyperphosphorylation of the STING-C206S mutant compared to STING WT. STING WT phosphorylation on Ser^366^ triggered by 2’3’-cGAMP was fully abrogated by HOCl treatment, while induction of STING-C206S phosphorylation was only slightly decreased (**Fig. 5C**). In the absence of 2’3’-cGAMP, HOCl did not alter basal STING WT phosphorylation, and only slightly diminished STING-C206S mutant phosphorylation (**Fig. 5D**). Altogether, these findings point to a role of Cys^206^ oxidation in the prevention of STING hyperactivation through the inhibition of STING Ser^366^ phosphorylation.

**Fig. 5.**
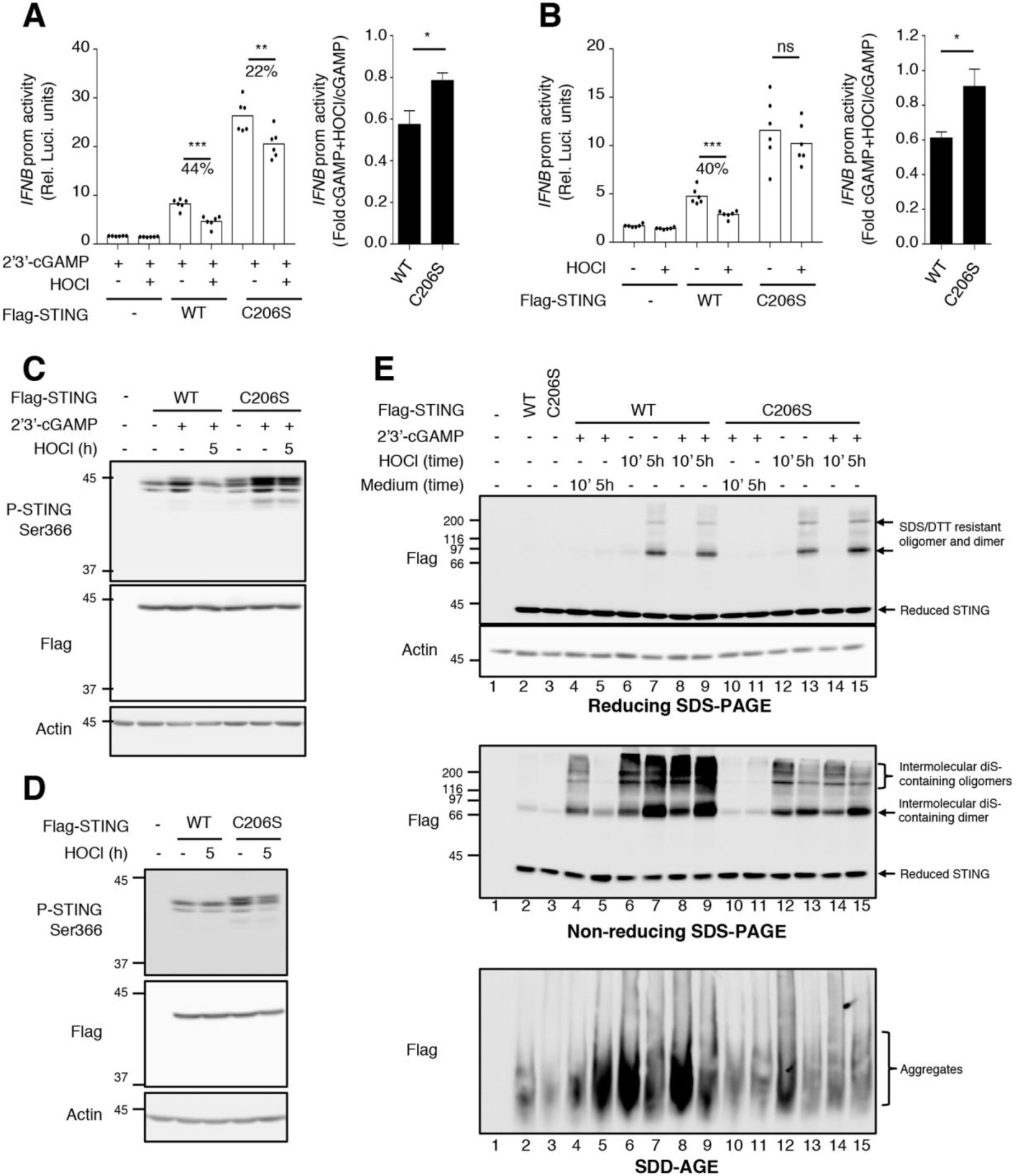
Cys^206^ oxidation dampens STING activity through inhibition of Ser^366^ phosphorylation and formation of inactive diS-containing polymers. **A-B**. HEK293T cells were transfected with the IFNβ promoter luciferase reporter construct together with the Flag-tagged WT or C206S STING-encoding plasmids or the corresponding control plasmid (empty). Cells were further transfected with 2’3’-cGAMP at 24µg/mL for 1h before treatment with 100µM hypochlorous acid (HOCl) for 23h (**A**) or were left untreated or stimulated with 100µM HOCl for 23h (**B**). Relative luciferase activities were measured. Mean +/- SEM, n=6, unpaired t-test. **C-E**. HEK293T cells (**C and D**) or A549 cells (**E**) were transfected with Flag-tagged WT or C206S STING-encoding plasmids or the corresponding control plasmid (empty). In **C**, cells were left untreated or stimulated with 100µM HOCl. In **D and E**, cells were further transfected with 2’3’-cGAMP at 8 (E) or 24 (D) µg/mL for 1h before treatment with 100µM HOCl for the indicated times. In **C-D**, Whole cell extracts were resolved by SDS-PAGE. In **E**, Whole cell extracts were resolved by reducing (with DTT) and non-reducing (without DTT) SDS-PAGE and by SDD-AGE. Proteins were detected by immunoblot using anti-actin, anti-Flag and anti-phospho-STING-Ser^366^ antibodies. Molecular weight (kDa) markers are indicated on the left side of the SDS-PAGE gels. The data are representative of three independent experiments.

We next assessed the state of polymerization of STING WT vs STING-C206S by non-reducing SDS-PAGE and SDD-AGE. Cells expressing Flag-tagged-STING WT or - STING-C206S were stimulated or not with 2’3’-cGAMP for 1h before further treatment with HOCl or the corresponding control medium for the indicated times (**Fig. 5E**). As expected based on observations made on endogenous STING (**Fig. 3C-E**), 2’3’-cGAMP stimulation triggered STING WT to form transient diS-containing oligomers of various sizes (**Fig. 5E, lanes 4-5 compared to lane 2, middle panel**) together with an accumulation of HMW aggregates (**Fig. 5E, lanes 4-5 compared to lane 2, lower panel**). The different diS-containing species were not detected with the STING C206S mutant and the HMW aggregates were strongly reduced (**Fig. 5E, lanes 10-11 compared to lanes 4-5, middle and lower panel**). Surprisingly, while HOCl alone did not activate, but rather inhibited STING, as measured by IFNβ luciferase reporter assay and Ser^366^ phosphorylation (**Fig. 5B and D**), it induced an accumulation of diS-containing dimeric and oligomeric species associated with transient formation of aggregates (**Fig. 5E, lanes 6-7 compared to lane 2, middle and lower panel**). It is noteworthy that the diS-containing species differ from those observed in response to 2’3’-cGAMP in that they are partially resistant to denaturing and reducing treatments (**Fig. 5E, lanes 6-7 compared to lanes 4-5, upper panel**). The profile was similar with STING-C206S although at a much-reduced level (**Fig. 5E, lanes 12-13 compared to lanes 6-7**). Finally, when cells were stimulated with 2’3’-cGAMP before HOCl treatment, STING WT and STING-C206S behaved similarly as with HOCl alone (**Fig. 5E, lanes 8-9 compared to lanes 6-7 and lanes 14-15 compared to lanes 12-13**). Overall, our data point to an inhibitory role of Cys^206^ ox-PTM associated with inhibition of Ser^366^ phosphorylation and formation of inactive diS-containing polymers.

## Discussion

The understanding of cell signaling regulation by redox processes has largely benefitted from the recent development of labeling techniques aimed at identifying protein subjected to reversible Cys ox-PTMs (*7, 15*). Although these techniques have allowed identification of numerous human proteins undergoing Cys ox-PTMs in vivo, the report of specific oxidation sites by MS is very often limited to proteins highly abundant and/or highly susceptible to oxidation (*13, 14, 67*). In most cases, identification of the oxidized residue by MS relied on the production of recombinant proteins and in vitro oxidation (*19-21*). In the present study, we successfully used Mal-PEG_2_-Bio-based labeling coupled to InsPEx workflow to identify novel Cys ox-PTM sites in vivo by LC-MS/MS (**Fig. 1**). Overall, we identified 2720 unique Cys ox-PTM sites encompassing 1473 proteins in U937 cells either unstimulated or subjected to treatment with diamide or HOCl (**Fig. 1D and Data File S3**).

The broad map of the identified proteins subjected to reversible Cys ox-PTM in vivo encompasses proteins with various activities, including those important for cellular signaling such as kinases, phosphatases, isomerases, transporters and receptors (**Fig. 2 and Data File S4**). Additionally, proteins subjected to reversible Cys ox-PTM exhibited various subcellular localizations, including cytoplasm, vacuoles/lysosomes compartments, ER, mitochondria and the nucleus (**Fig. 2**). Of interest to us, we identified numerous proteins potentially involved in virus-host interaction. Several BPs enriched in proteins harboring Cys ox-PTM were related to translation and intracellular localization regulation (intracellular transport, establishment of protein localization, protein transport and nuclear transport) (**Fig. 2**). Functional clusters analysis highlighted the importance of translation frameshifting termination, and transport multivesicular localization. The translational and intracellular transport functions found regulated by oxidative stress are relevant to host-virus interaction because of their involvement in cell signaling regulation, but also because they are key for proper virus replication and assembly (*68*). KEGG pathways analysis revealed selective enrichment of Cys ox-PTMs in RNA transport pathway, which include several components of the nuclear pore function, namely nucleoporins and exportins (**Fig. 2F, fig. S4B and Data File S5**). Transport in and out of the nucleus is particularly important for viruses with steps of their replication cycle occurring in the nucleus, such as influenza virus (*69, 70*). Additionally, numerous viruses hijack the nuclear pore through NUPs to regulate trafficking of viral proteins, genomes and even capsids into and out of the nucleus, thus promoting their replication (*71*). Previous reports have shown that oxidative stress regulates export from the nucleus through post-translational modifications of NUPs (*72, 73*). The reversible Cys ox-PTMs sites in NUPs and exportins identified in this study likely represents novel regulatory mechanisms of nuclear transport. Several BPs enriched in proteins harboring Cys ox-PTMs were also related to the immune response (immune system processes and leukocyte activation) (**Fig. 2**). For instance, we identified several novel Cys ox-PTM sites of components of the major histocompatibility complex I (MHC-I) upon oxidant treatment: HLA-A Cys188, Cys227, and Cys363; HLA-C Cys188 and Cys227; HLA-DRA Cys188. Above all, analyzes of the KEGG pathways revealed numerous proteins specifically involved in the interaction of the host with RNA (influenza) and DNA viruses (Herpes simplex and HIV) subjected to Cys ox-PTMs (**Fig. 2F**). In particular, this study provides the first direct demonstration that the adaptor STING, which is key to the IFN antiviral and proinflammatory response triggered following DNA virus sensing, is subjected to reversible Cys ox-PTMs.

Previous reports have documented contradictory evidence of redox regulation of the STING pathway. In some studies, modulation of the nuclear factor erythroid 2-related factor 2 (Nrf2), which is essential for the production of antioxidants and protection of cells against oxidative stress, decreased 2’3’-cGAMP or DNA virus-induced STING-dependent responses, suggesting a positive regulation by ROS (*74-76*). In contrast, ROS appear to inhibit PPARγ-mediated activation of STING-dependent IFN expression (*77*). Inhibition of ectopically expressed STING activity was also observed upon induction of ROS by the mitochondrial complex I inhibitor rotenone (*60*). Certainly, several reports have provided evidence pointing to a role of Cys residues in the redox regulation of STING in that they showed formation of diS-containing dimers and aggregates sensitive to reducing agents upon ligand stimulation (*58, 60, 61, 78-80*). However, the identity of Cys subjected to ox-PTM and their role remained to be elucidated.

Here, we demonstrate oxidation of Cys^148^ under basal conditions and upon treatment with HOCl, while oxidation of Cys^206^ was inducible by HOCl and by the natural 2’3’-cGAMP ligand (**Fig. 3 and fig. S5**). Previous reports have shown that STING-C148A and -C148S mutants exhibit impaired capacity to transit to the golgi and activate the IFNβ promoter, thereby supporting that Cys^148^ oxidation is necessary for STING activation (*60, 61*). Although formation of diS-bound dimer upon H2O2 and 2’3’-cGAMP stimulation was also impaired, the oxidation of Cys^148^ was not directly demonstrated (*60, 61*). Despite the fact that none of the available human ligand-bound STING structures allows to see the linker region containing Cys^148^ in electronic density maps (*29, 61-65, 81-91*), it was proposed that Cys^148^ is engaged in a ligand-inducible diS bond that stabilizes the uncrossed linker regions in the STING polymer (*60, 61*). Our data, however, argues in favor of a different model in which Cys^148^ oxidation is already present at basal levels. In agreement with previous reports (*60, 61*), we did not detect dimers or oligomers containing intermolecular diS at basal levels, inferring that Cys^148^ is not engaged in intermolecular diS bond, but rather subjected to another type of ox-PTM that remains to be identified. We evaluated the presence of intramolecular diS bond by non-reducing SDS-PAGE (**fig. S6**). Although our results suggest the existence of a previously unrecognized intramolecular diS in ectopically ex pressed STING, it appears to be independent of Cys^148^. Thus, this study adds to previous reports to support a model in which Cys^148^ oxidation is essential for STING activity.

The observation that the 2’3’-cGAMP stimulation of the STING-C206S mutant leads to hyperactivation of STING together with the demonstration that HOCl inhibits STING activity in a Cys^206^-dependent manner (**Fig. 4F and 5A-B**) support an inhibitory role of the oxidation of Cys^206^ induced by 2’3’-cGAMP. This inhibition is associated with decreased Ser^366^ phosphorylation and formation of diS-containing polymers. Our finding that impairment of Cys^206^ oxidation through mutation into Ser leads to constitutive activity that is further hyperactivated in response to 2’3’-cGAMP is in agreement with a previous study reporting a patient with a gain-of-function mutation of Cys^206^ (pC206Y) associated with the STING-associated vasculopathy with onset in infancy (SAVI) type I interferonopathy (*92*). The gain-of-function phenotype was recapitulated by mutation of Cys^206^ into Phe, Leu and Ser (*92*). In this same study, patients with gain-of-function mutations of Arg^281^ (pR281Q) or (pR284G) that are spatially located close to Cys^206^ (**Fig. 4B-C**) were also identified. Arg residue are known to lower the pKa of Cys residue, thereby rendering ox-PTM possible (*93, 94*). The close proximity of Cys^206^ and Arg residues in the STING structure, together with the observation that Arg^281^ and Arg^284^ mutations mimic the gain-of-function observed with mutations of Cys^206^ that abolish ox-PTM, might suggest Arg^281^ and Arg^284^ are essential for Cys^206^ to be oxidized. However, further characterization of the STING-R284S mutant that recapitulates the gain-of-function phenotype, as did R284G, R284H and R284K mutations, revealed constitutive aggregation and formation of perinuclear puncta that were not further enhanced by 2’3’-cGAMP stimulation (*60, 61*) (*92*). This differs from our observation that STING-C206S constitutively active phenotype can by hyperactivated by 2’3’-cGAMP and is not associated with constitutive polymerization or localization in perinuclear puncta, but rather with increased phosphorylation (**Fig. 4F and 5**). Altogether, this argues against a role of Arg^284^ in Cys^206^ oxidation.

Predictions from the currently available cGAMP-bound unphosphorylated STING dimer structure do not show direct engagement of Cys^206^ in diS bond. This could suggest that Cys^206^ is engaged in interdimer diS bond or in an interaction with a yet to be identified protein partner rather than in the formation of STING dimers. Those diS bonds would not be visible in the structures of the dimer. Alternatively, this could also suggest that Cys^206^ is subjected to an ox-PTM distinct from diS but critical for the latter to form between other residues that were not detectable using our MS workflow. Ultimately, the determination of the exact oxidative modification of Cys^148^ and Cys^206^ would benefit from the development of novel dynamic models of STING structure taking into account not only the binding of the ligand but also the conformational changes induced by phosphorylation and oxidation. Meanwhile, in silico molecular modeling of the structureof ligand-bound STING containing sulfenylated Cys^206^ allowed us to get insight into the mechanism of STING inhibition. Based on the steric constraints imposed by the sulfenylation of Cys^206^ (**Fig. 4 B-C**), it is very likely that once oxidized, STING must undergo a change of conformation (*66*). Although we cannot exclude other mechanisms, it is reasonable to propose that this new conformation is not favorable for the binding of the TBK1 kinase or for TBK1 to access Ser^366^, thus explaining the Cys^206^-dependent inhibition of 2’3’-cGAMP-induced Ser^366^ phosphorylation by HOCl (**Fig. 5**). In agreement with previous reports (*58, 60, 61, 78-80*), we observed that oxidized STING induced by 2’3’-cGAMP is associated with the formation of polymers of different sizes containing diS (**Fig. 3 and 5**). Here, we show that mutation of Cys^206^ that results in STING hyperactivation by 2’3’-cGAMP leads to a loss of these polymers. In addition, HOCl inhibits STING activity while triggering the formation of the diS-containing polymers in a Cys^206^-dependent manner (**Fig. 5**). Taken together these observations suggest that the diS-containing polymers induced by 2’3’-cGAMP constitute inactive STING species. We also found that oxidative stress alone is sufficient to trigger the formation of inactive diS-containing oligomers and aggregates (**Fig. 5E**). These species are most likely distinct from the ligand-induced oligomers previously reported to be indispensable for STING binding to TBK1 and subsequent phosphorylation (*85, 88, 95*). Mutations which disrupted the formation of oligomers detected by gel filtration chromatography or native-PAGE, methods which do not assess the diS content, correlated with the inhibition of STING signaling in response to 2’3’-cGAMP (*85, 95*). These species also differ from those detected in our study in that they were found to contain phosphorylated Ser^366^ (*95*), whereas we did not detect phosphorylation of oligomers containing diS (**Fig. 3D**). Together, these observations support that distinct polymeric STING species that correspond to the active and inactive states can be detected after stimulation with 2’3’-cGAMP. The distinction of the state of activity cannot be determined solely with the size of the polymer, but must include the study of the content in diS dependent on Cys^206^.

Collectively, our data establish a novel model of STING regulation by ox-PTMs (**fig. S8**). First, STING Cys^148^ is oxidized independently of ligand binding or induction of oxidative stress. According to previous reports, this oxidation state is essential for STING to be activated by 2’3’-cGAMP (*61*). Next, binding of 2’3’-cGAMP triggers Ser^366^ phosphorylation by TBK1 and activation of STING leading to expression of IFNβ. Further oxidation of Cys^206^ in response to 2’3’-cGAMP imposes a conformational change that inhibits Ser^366^ phosphorylation and provokes the formation of inactive diS-containing polymers to prevent STING hyperactivation. This study paves the way to the design of novel STING agonists or antagonists for treatment of autoinflammatory disorders and development of immunotherapies and vaccines.

## Materials and Methods

### Cell culture

The monocytic cell line U937 (obtained from Dr. M. Servant, Université de Montréal, Quebec, Canada) was cultured in RPMI medium (GIBCO) supplemented with 1% L-glutamine (GIBCO), 1% HEPES (GIBCO) and 10% Fetalclone III (FCl-III, Hyclone). The A549 and human embryonic kidney (HEK) 293T cell lines (American Type Culture Collection, ATCC) were respectively grown in F12 and DMEM medium (GIBCO), both supplemented with 1% L-glutamine and 10% FCl-III. No antibiotics were used for cell culture. The MycoAlert Mycoplasma Detection Kit (Lonza) was used to monitor mycoplasma contamination on a regular basis.

### Plasmids

The pcDNA3.1 Flag-STING plasmid, encoding for the human protein, was obtained from Dr. D. Lamarre, Université de Montréal, Montréal, Canada. The pcDNA3.1 Flag-STING-C206S construct was generated using the Quikchange Lightning Site-Directed Mutagenesis kit from Agilent (#210518) with the following primer (Invitrogen): 5’-TTGAAATTCCCTTTTTCACTCACTGCAGAGATCTCAGC-3’. Mutation and STING sequence was validated by Sanger sequencing at the Génome Québec Innovation Centre (McGill University, Montréal, QC).

### Plasmid transfection

Plasmid transfection in HEK293T cells was achieved using the calcium phosphate co-precipitation method. A total of 2.5µg of DNA was transfected per 24 well for 24h. For A549, the TransIT-LT1 transfection reagent (Mirus) was used to transfect a total of 3µg of DNA per 35mm plate of cells at a confluency of 70%, with a TransIT/DNA ratio of 1:2 as per manufacturer instructions.

### Oxidants preparation

Diamide (Sigma Aldrich) was prepared from a 0.5M stock solution in ddPBS (GIBCO) by dilution in DMEM without phenol red (DMEM w/o PR, GIBCO) containing 1% L-glutamine and 2% FCI-III. Hypochlorous acid (HOCl) was prepared by 1/5 dilution of NaOCl solution with available chlorine 4.00-4.99 % (Sigma Aldrich) in ddH2O containing 0.1M NaOH. The pH was adjusted to 11 with 12.1M HCl (BioShop). HOCl concentration was quantified by measure of absorbance at 292nm with a molar absorption coefficient of 350M-1cm-1. The final solution at the desired concentration was obtained by dilution in DMEM w/o PR supplemented with 2% FCI-III and 1% L-glutamine.

### Cells treatment with oxidants

For U937 cells treatment, cells were plated at a concentration of 10^6^ cells/mL in RPMI supplemented with 1% L-glutamine, 1% HEPES and 10% FCl-III. Sixteen hours before stimulation, the medium was replaced by DMEM w/o PR containing 2% FCI-III and 1% L-glutamine. At the time of stimulation, the medium was replaced by DMEM w/o PR containing 1% L-glutamine, 2% FCI-III and 0.5mM diamide or 100µM HOCl. Treatments were pursued for the indicated times. For HEK293T and A549 cells transfected with STING-encoding plasmids, cells were treated with HOCl at 100µM for the indicated times in the corresponding medium containing 2% FCI-III and 1% L-glutamine. Where indicated, cells were transfected with 2’3’-cGAMP as detailed below 1h before HOCl treatment.

### 2’3’-cGAMP transfection

In U937 cells, infusion of 2’3’-cGAMP (Invivogen) was carried out by digitonin permeabilization as previously described in (*96*). Briefly, cells at 90% confluency were incubated at 37°C for 30min in a buffer containing 50mM Hepes pH 7.0, 100mM KCl (Sigma Aldrich), 3mM MgCl2 (Sigma Aldrich), 85mM sucrose (Sigma Aldrich), 0.2% (m/v) BSA (Sigma Aldrich), 0.1mM DTT, 10µg/mL digitonin (Sigma Aldrich), 1mM ATP (Sigma Aldrich), 0.1mM GTP (Sigma Aldrich) with or without 2’3’-cGAMP at a final concentration of 24µg/mL. After 30min, the buffer was replaced with RPMI containing 10% FCl-III, 1% L-glutamine, 1% HEPES. Stimulations were pursued for the indicated times. 2’3’-cGAMP transfection in HEK293T and A549 cells was achieved using the Lipofectamine 2000 transfection reagent (Invitrogen). Briefly, 2’3’-cGAMP at 8 or 24µg/mL was used to transfect cells at 90% confluency using a Lipofectamine ratio of 1:1 in medium containing 2% FCl-III. For transfections longer than 6h, medium was replaced by fresh medium supplemented with 2% FCl-III.

### Bioswitch labeling of Cys ox-PTMs

Unless otherwise stated, all steps were performed on ice and in the dark. At time of harvesting, cells were immediately transferred to pre-chilled tubes and washed in cold ddPBS before trichloroacetic acid (TCA, Sigma Aldrich)/acetone (Fisher chemical) precipitation (*11*). After TCA/acetone addition, samples were sonicated (3×15 pulses, 70%, UP50H sonicator, Hielscher) with rounds on ice. Samples were stored at −80°C for at least 2h before further processing. The bioswitch labeling procedure was performed according to the protocols described in (*11, 45*) with modifications using N-ethylmaleimide (NEM, Sigma Aldrich) to alkylate the reduced thiols, tris(2-carboxyethyl)phosphine (TCEP, EMD Millipore Sigma) to reduce the oxidized thiols and EZ-Link Maleimide-PEG_2_-Biotin (Mal-PEG_2_-Bio, Thermofisher) to label the newly reduced thiols (**Fig. 1A**). Briefly, 12mg of TCA/acetone precipitated proteins were used for alkylation with 3mM NEM in 0.5M Tris, 1% SDS (BioShop) pH 7.3 for 30min at room temperature (RT). Next, reduction was performed using 4mM TCEP in 0.5M Tris, 1% SDS pH 7.0 for 1h at RT followed by labeling with 0.25mM Mal-PEG_2_-Bio for 2h at RT.

### In-solution trypsin digestion of Mal-PEG_2_-Bio-labeled proteins

In the In-solution trypsinization with Peptide Extraction (InsPEx) method (**Fig. 1B**), Mal-PEG_2_-Bio-labeled proteins were precipitated with acetone. Pellets were resuspended in 0.5M Tris containing 1% SDS pH 7.0 before quantification using the BCA (Pierce Thermoscientific) method. Ten mg of proteins were acetone precipitated and pellets were resuspended in 50mM Tris-HCl pH 7.0, 8M Urea (Sigma Aldrich), 20mM 1,4-Dithiothreitol (DTT, Sigma Aldrich) before incubation at 60°C for 1h. Next, samples were diluted with 50mM Tris-HCl pH 7.0 to decrease urea concentration to 1M. Trypsin was added to a final protease:protein ratio of 1:30 (w/w) and digestion was performed ON at 37°C. To maximize the digestion, additional Trypsin was added after 4h of incubation to a final protease:protein ratio of 1:30 (w/w).

### Liquid-solid phase extraction of trypsin-digested peptides

Trypsin activity was stopped by addition of 0.1% trifluoroacetic acid (TFA, Sigma-Aldrich), pH 1.0 to the digestion mix described above before centrifugation at 20000g for 20min at RT. The supernatant containing the digested peptides was transferred to a new tube and the pellet was further extracted with a solution of 0.1% TFA to make sure all digested peptides were collected. Supernatants of both extractions were pooled and used for the next step (**Fig. 1B**). Peptides were desalted and extracted, according to the protocol described in (*97*), with modifications detailed in (*98*). In brief, Sep-Pak tC18 solid-phase extraction cartridges (Waters) were sequentially equilibrated with 1) 100% acetonitrile (Sigma Aldrich), 2) 0.5% acetic acid (Sigma Aldrich) and 50% acetonitrile, and 3) 0.1% TFA. Supernatants containing the digested peptides were loaded on the cartridges and sequentially washed with 0.1% TFA and with 0.5% acetic acid. Peptides were eluted into a clean tube with 1mL of 0.5% acetic acid and 80% acetonitrile and distributed into multiple tubes containing 150µL before complete evaporation using a speed vacuum concentrator (Savant SVC 100H).

### NeutrAvidin pull down of Mal-PEG_2_-Bio-labeled peptides

The dried peptide pellets described above were resuspended in HENS buffer composed of 250mM HEPES (BioShop) pH 7.7, 1mM EDTA (BioShop), 0.1mM Neocuproine (Sigma Aldrich) and 1% SDS before addition of 2 volumes of neutralization buffer containing 20mM HEPES pH 7.7, 100mM NaCl (BioShop), 1mM EDTA and 0.5% (v/v) Triton X-100 (BioShop). Incubation with 200µL (50% slurry) NeutrAvidin agarose resin (ThermoFisher, Pierce) was performed overnight (O/N) at 4°C. After 5 washes with a solution of 20mM HEPES pH 7.7, 600mM NaCl, 1mM EDTA and 0.5% Triton X-100, 2 washes with ddPBS and a final wash with UltraPure Distilled Water (Invitrogen), peptides were eluted with 8M Guanidine/HCl (Sigma Aldrich) pH 1.5.

### LC-MS/MS mass spectrometry

Before LC-MS/MS analysis, the pH of eluted peptides was corrected to 7.4 by addition of 1M Tris/HCl, pH 7.4. LC-MS/MS analysis was performed on an Orbitrap Fusion Lumos Tribrid mass spectrometer (Thermo Scientific) operated with Xcalibur (version 4.0.21.10) and coupled to a Thermo Scientific Easy-nLC (nanoflow Liquid Chromatography) 1200 system at the Southern Alberta Mass Spectrometry (SAMS) facility of the University of Calgary (Alberta, Canada). Tryptic peptides were loaded onto a C18 trap (2 cm × 75 um; ThermoScientific PN164946) at a flow rate of 2ul/min. Peptides were then eluted and separated using an Easy Spray Column (50 cm × 75 um; ThermoScientific PN ES803) and a 45 min gradient from 5 to 40% (5% to 28% in 40 min followed by an increase to 40% B in 5 min) of solvent B (0.1% formic acid in 80% LC-MS grade acetonitrile) at a flow rate of 0.3 μL/min. Solvent A was composed of 0.1% formic acid and 3% acetonitrile in LC-MS grade water. Peptides were then electrosprayed using 2.3 kV voltage into the ion transfer tube (300°C) of the Orbitrap Lumos operating in positive mode. The Orbitrap first performed a full MS scan at a resolution of 120000 FWHM to detect the precursor ion having a m/z between 375 and 1575 and a +2 to +7 charge. The Orbitrap AGC (Auto Gain control) and the maximum injection time were set at 4e5 and 50 ms, respectively. The Orbitrap was operated using the top speed mode with a 3 sec cycle time for precursor selection. The most intense precursor ions presenting a peptidic isotopic profile and having an intensity threshold of at least 5000 were isolated using the quadruple and fragmented with HCD (30% collision energy) in the ion routing multipole. The fragment ions (MS2) were analyzed in the ion trap at a rapid scan rate. Dynamic exclusion was enabled for 30 sec to avoid the acquisition of same precursor ion having a similar m/z (plus or minus 10 ppm).

### LC-MS/MS data analysis

The Lumos data files were converted into Mascot Generic Format (MGF) using RawConverter (v1.1.0.18; The Scripps Research Institute) operating in a data dependent mode. Monoisotopic precursors having a charge state of +2 to +7 were selected for conversion. This mgf file was used to search a database (Homo sapiens, NCBI; 99742 entries) using Mascot algorithm (Matrix Sciences; version 2.4), assuming the digestion enzyme trypsin at a maximum of 1 missed cleavage. Mascot was searched with a fragment ion mass tolerance of 0.60 Da and a parent ion tolerance of 10.0 PPM. Oxidation of methionine (Met+15.99Da), alkylation of Cys with NEM (Cys+125.05Da) and modification of Cys with Mal-PEG_2_-Bio (Cys+525.23Da, **Fig. 1C**) were specified in the Mascot search parameters as variable modifications. The Mascot data files were imported into Scaffold (v4.8.4, Proteome Software Inc., Portland, OR) for comparison of different samples based on their mass spectral counting. Scaffold was used to validate MS/MS based peptide and protein identifications. For label-free quantification (LFQ), only peptides containing at least 1 Cys ox-PTMs and detected in at least two out of the three biological replicates were considered for spectral counting. Scaffold algorithms were used to perform standard cutoffs of significance for peptide ion signal peak intensity-based quantification (*52, 99*). Peptide identifications were accepted if they could be established at greater than 95.0% probability by the Peptide Prophet algorithm (*100*) with Scaffold delta-mass correction and a false discovery rate (FDR) <1%. Protein identification was accepted if it could be established at greater than 95.0% probability and contained at least 1 identified oxidized peptide. Protein probabilities were assigned by the Protein Prophet algorithm (*101*). Proteins that contained similar peptides and could not be differentiated based on MS/MS analysis alone were grouped to satisfy the principles of parsimony.

### Gene Ontology and KEGG pathways analysis

Identified proteins containing Mal-PEG_2_-Bio-labelled Cys were subjected to Gene Ontology (GO) enrichment analysis using the Cytoscape software (version 3.6.0) with the BiNGO plugin (*53, 54*), which compare the number of labeled proteins against the total proteins known to belong to each GO term. To assess the enrichment of a GO term, the hypergeometric test was used. The Benjamini and Hochberg procedure was applied to control the FDR and correct the p values (*102*). Adjusted p-value <0.05 were considered significant. Networks with overrepresented GO categories after correction were mapped. For pathway analyses, the ClueGO plugin was used (*55*) with the analysis mode “functions” to list the pathways enriched in Mal-PEG_2_-Bio-labelled Cys within the Kyoto Encyclopedia of Genes and Genomes(KEGG) database. Statistical analyses were done using a hypergeometric test, with the Benjamini and Hochberg procedure for p-value correction. ClueGO layout was used to visualize enriched pathways.

### Preparation of Whole Cell Extracts (WCE)

Samples were kept on ice during all steps. Cells were harvested in cold ddPBS, transferred into 1.5mL tubes and centrifuged at 16200g, 4°C for 20sec. Pellets were resuspended in 70µL RIPA lysis buffer constituted of 50mM Tris pH 7.5, 150mM NaCl, 2mM EDTA, 0.1% SDS, 1% Triton and 1% Sodium Deoxycolate completed with 10µg/mL leupeptin, 10µg/mL aprotinin, 10µg/mL pepstatin, 30mM NaF, 1mM activated Na3VO4, 1mM p-nitrophenyl phosphate and 10mM β-glycerophosphate. For reducing vs non-reducing SDS-PAGE experiments, RIPA buffer was also supplemented with 100mM NEM. After 20min incubation for in lysis buffer, samples were sonicated at 40% with 3×15 pulses of 0.8sec with rounds on ice in between. Samples were centrifuged at 800g at 4°C for 10min. The supernatants were kept and quantified using the DC method (DC Protein Assay Kit, Bio-Rad).

### Semi-Denaturing Detergent Agarose Gel Electrophoresis (SDD-AGE)

Fresh WCE were resolved by SDD-AGE following the protocol detailed in (*103 9356*) with the modifications reported in (*104*). Briefly, thirty µg of WCE completed with one-third volume of 4X loading dye (20% glycerol, 8% SDS and bromophenol blue in 2x TAE buffer) were incubated for 5min at RT before resolution in a vertical 1.5% agarose gel. The running buffer consisted of TAE 1X supplemented with 0.1% SDS. To transfer the proteins to a nitrocellulose membrane, a liquid transfer was performed overnight at 4°C at 20V constant using transfer buffer containing 25mM Tris-Base, 192mM glycine, 0.1% SDS and 10% EtOH. To better exposed the epitopes, membranes were treated with 0.5% glutaraldehyde (LifeSensors) in dPBS for 20 min before immunoblots (*105 9872*).

### Non-reducing/reducing SDS-PAGE

SDS-PAGE electrophoresis was performed as detailed in (*106*) using 12% acrylamide/bis-acrylamide (37.5:1, Bioshop) resolution gel and a stacking gel of 3% acrylamide/bis-acrylamide (37.5:1). Fresh WCE (30 or 50µg) were completed with one-fourth volume of 5X loading buffer (125 mM Tris-HCl pH 6.8 (RT), 10% SDS, 20% glycerol, 0.0005% bromophenol blue). For the reducing conditions, 250mM DTT was added to the loading buffer. Samples were incubated for 5min at 100°C. Heating was quenched by a 2min incubation on ice. Both the stacking and resolution gels were transferred onto nitrocellulose using a buffer supplemented with 0.036% SDS and 10% Ethanol.

### Immunoblot analysis

Following protein transfer, nitrocellulose membranes were probed with protein-specific primary antibodies diluted in PBS containing 5% Milk or 5% BSA as follows: anti-actin Cat #A5441 from Sigma Aldrich; anti-tubulin Cat #5286 from Santa Cruz; anti-Flag M2 Cat #SLBF6631V from Sigma Aldrich; anti-Phospho-STING (Ser^366^) Cat #19781 from Cell signaling; anti-STING Cat #13647 from Cell signaling; anti-Phospho-TBK1 (Ser^172^) Cat #5483 from Cell signaling; anti-TBK1/IKKε Cat #IMG270A from Imgenex. Membranes were washed using dPBS containing 0.5% Tween (PBS-T) before incubation with horseradish peroxidase (HRP)-conjugated secondary antibodies (Seracare or Jackson Immunoresearch Laboratories). After final wa shes, enhanced chemiluminescence using the Western Lightning Chemiluminescence Reagent Plus (Perkin-Elmer Life Sciences) was performed to identify immunoreactive bands detected using a LAS4000mini CCD camera apparatus (GE healthcare).

### Immunofluorescence microscopy

Transfected HEK293T were seeded in Poly-L-Lysine coated 96-well clear bottom plates (Corning Incorporated; plate number 3882 half-area microplates) before fixation with 3.7% formaldehyde for 15min at 37°C and permeabilization with 0.5% Triton X-100 in ddPBS containing 0.5mM CaCl2, 1mM MgCl2 (PBS/Ca/Mg) for 20min at RT. After blocking with 2% BSA, 0.2% Tween-20 in PBS/Ca/Mg for 20min at RT, protein were immunodetected by incubation overnight at 4°C with the Anti-DYKDDDDK Tag (D6W5B) Rabbit mAb (Cell Sign #14793; recognizes the Flag epitope) followed by Donkey anti-Rabbit, Alexa Fluor 647 (Invitrogen A-31573) secondary antibody for 90min at RT. Nuclei were counterstained with 4’,6’-Diamidino-2-phenylindole (DAPI; #D9564; Sigma) diluted 1/1500 in PBS/Ca/Mg for 5min. Plates were stored in the dark at 4°C prior to imaging using a 40x objective mounted on an Operetta High-Content Imaging System (Perkin Elmer) controlled by Harmony 4.1 software.

### Structure analysis

The available STING structures in the protein data bank (PDB) database (*107*) were downloaded and visualized using PyMOL (Schrödinger, Inc. version 2.2.3). The PyTM plugin was used to sulfenyl (S-OH) on Cys^206^. Sulfenylation was used given that it is the first ox-PTMs happening before subsequent glutathionylation or disulfide bond formation (*7, 108*). The APBS Electrostatics plugin was used to visualize the electrostatic surfaces (*109, 110*). The CEalign plugin was used to perform superposition of the structures and the computation of the difference of the root-mean-square deviation of atomic positions (RMSD) (*111*).

### IFNβ promoter luciferase gene reporter assay

HEK293T cells were cotransfected with the IFNβ-pGL3 firefly (*112*), the pRL-null renilla (Promega, internal control) luciferase reporter plasmids together with the STING encoding plasmid. At 24h, cells were stimulated or not with 100µM HOCl for 24h. Where indicated cells were transfected with or without 2’3’-cGAMP at 8 or 24µg/mL 1h before treatment with HOCl. The cells were collected and assayed for reporter gene activities using the Dual-luciferase reporter assay system (Promega). The firefly/renilla luciferase ratios were calculated to determine luciferase activities. Statistical analyses were performed using the Prism 8 software (GraphPad) using the indicated tests. Statistical significance was assessed using the following p values: p < 0.05 (*), p <0.01 (**), p <0.001 (***) or p <0.0001 (****). p values <0.05 were considered significant.

## Supplementary Materials

Data file S1. Mal-PEG_2_-Bio-modified peptides found in the pilot in-gel trypsinization approach in THP1 cells upon diamide treatment.

Data file S2. Mal-PEG_2_-Bio-modified peptides found using the InsPEx approach in U937 cells upon diamide treatment.

Data file S3. Biological triplicates of Mal-PEG_2_-Bio-modified peptides found with the InsPEx bioswitch approach at basal levels and upon diamide and HOCl stimulations.

Data file S4. List of all Gene Ontology (GO) terms that are enriched in Cys ox-PTM at basal levels and upon diamide/HOCl stimulations. Biological processes, molecular functions and cell component are listed in different sheets.

Data file S5. List of all KEGG pathways found enriched in oxidized proteins at basal levels and upon diamide/HOCl stimulations.

## Acknowledgments

We thank Dr. M. Servant (Université de Montréal, Quebec, Canada) for providing U937 cell line, Dr. K. Gee (Queens University, Ontario, Canada) for THP-1 cells and Dr. D. Lamarre (Université de Montréal, Quebec, Canada) for the STING encoding plasmid. We thank Dr. L. Brechenmacher at the Southern Alberta Mass Spectrometry (SAMS) facility of the University of Calgary (Alberta, Canada) for expert technical assistance. We also thank Dr. P. Arthur (University of Western-Australia, Australia) for help with the bioswitch labeling protocol and Dr. E. Lecuyer and X. Wang for help with immunofluorescence staining.

## Author contributions

NZC and NG designed the experiments. AF and SC optimized the bioswitch labeling technique. NZC and EC performed the experiments. NZC and NG analyzed the data and wrote the manuscript.

## Data and materials availability

Proteome wide mass spectrometry data will be available in the PRIDE database, accession number [xxxxxxx].

## Supplementary Materials

### Supplementary Materials and Methods

#### Plasmids

The pcDNA3.1 Flag-STING-C148S was generated using the Quikchange Lightning Site-Directed Mutagenesis kit from Agilent (#210518) with the 5’-TCAGGCACCCC**<u>ACT</u>**GTCCAATGGGAGG-3’ primer. Sanger sequencing at the Génome Québec Innovation Centre (McGill University, Montréal, QC) was used to validate the mutation and STING sequence.

#### NeutrAvidin pull-down of Mal-PEG_2_-Bio-labeled proteins followed by in-gel trypsin digestion (in-gel trypsinization approach, **fig. S1B**)

THP-1 cells (obtained from Dr. K. Gee, Queens University, Ontario, Canada) were cultured in RPMI medium (GIBCO) supplemented with 1% L-glutamine (GIBCO), 1% HEPES (GIBCO) and 10% Fetalclone III (FCl-III, Hyclone). Cells were subjected to stimulation with 0.5mM diamide for 20min. Proteins were subjected to Mal-PEG_2_-Bio bioswitch labeling as described in Material and Methods section. Mal-PEG_2_-Bio-labeled proteins were precipitated with acetone. Pellets were resuspended in HENS buffer and quantified by the BCA method. One milligram of protein was taken before addition of 2 volumes of neutralization buffer. Pull down with NeutrAvidin agarose resin was performed as described in the Material and Methods section with modifications of the elution step. Proteins were eluted with 2X loading buffer composed of 50 mM Tris/HCl pH 6.8, 4% SDS, 8% glycerol (BioShop), 0.0005% bromophenol blue (Fisher Scientific) supplemented with 1.44 M β-mercaptoethanol (BME, Sigma Aldrich). Samples were heated at 100 °C for 10min with 2 vortexing rounds to harvest all complexes. Samples were loaded on 1mm SDS-PAGE gels. In order to obtain a single band containing all the proteins, the migration was stopped once the samples reached around 5mm distance into the resolution gel. After staining with Coomassie Brilliant Blue R-250 Dye (BIO-RAD), the single band was cut and chopped into small pieces. In-gel trypsin (Pierce™ Trypsin Protease, MS Grade) digestion was performed as recommended by the manufacturer.

#### Analysis of functional clusters in GO networks enriched in Cys ox-PTMs

Networks with overrepresented GO categories identified as described in the Material and Methods section and represented in (**Fig. 2A-C**) were subjected to further analysis using the greedy community-structure detection algorithm (GLay), incorporated in Cytoscape to identify functional clusters within the networks (*113*).

## Supplementary Figure legends

**Figure S1.**
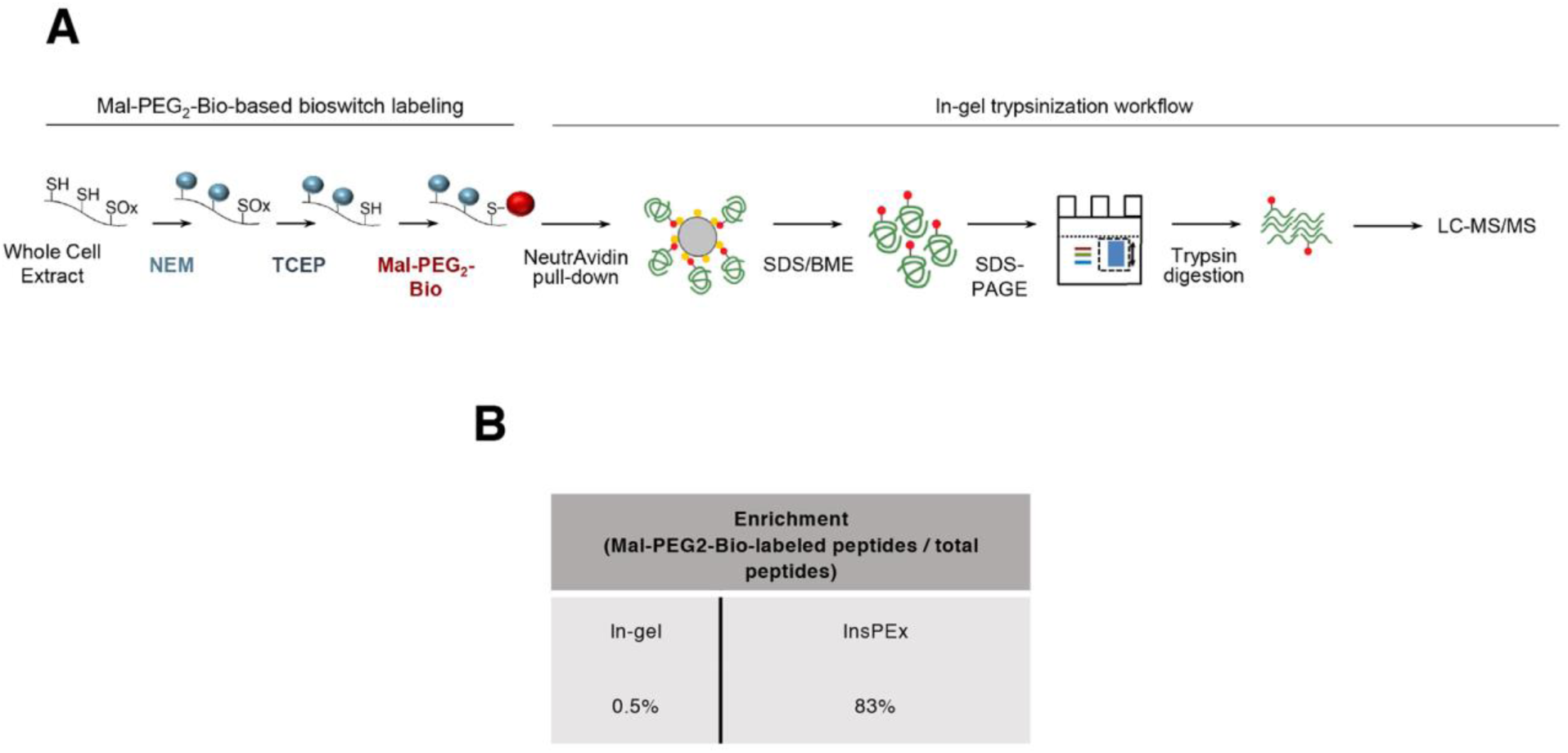
Workflow of the bioswitch labeling strategy coupled to in gel trypsin digestion for identification of Cys ox-PTMs by MS in vivo. *Supplemental figure related to Fig. 1*. **A-B**. Schematic of the workflow used to label reversible Cys ox-PTMs with the Mal-PEG_2_-Bio bioswitch method (**A**) followed by In-gel trypsinization (**B**). In (**A**), Cys-SH residues were specifically alkylated with N-ethylmaleimide (NEM). Reversibly oxidized Cys (SOx) were then reduced using tris(2-carboxyethyl)phosphine (TCEP) before labeling with EZ-Link Maleimide-PEG_2_-Biotin (Mal-PEG_2_-Bio). In (**B**), Proteins from (**A**) were pulled down using NeutrAvidin agarose beads before loading the eluate on SDS-PAGE gel. Peptides were generated by in-gel trypsin digestion before LC-MS/MS analysis. **C**. Table showing the Enrichment (percentage of Mal-PEG_2_-Bio labelled peptides amongst total peptides) obtained with the InsPEx (Fig. 1A-B) and the In-gel trypsinization strategies.

**Figure S2.**
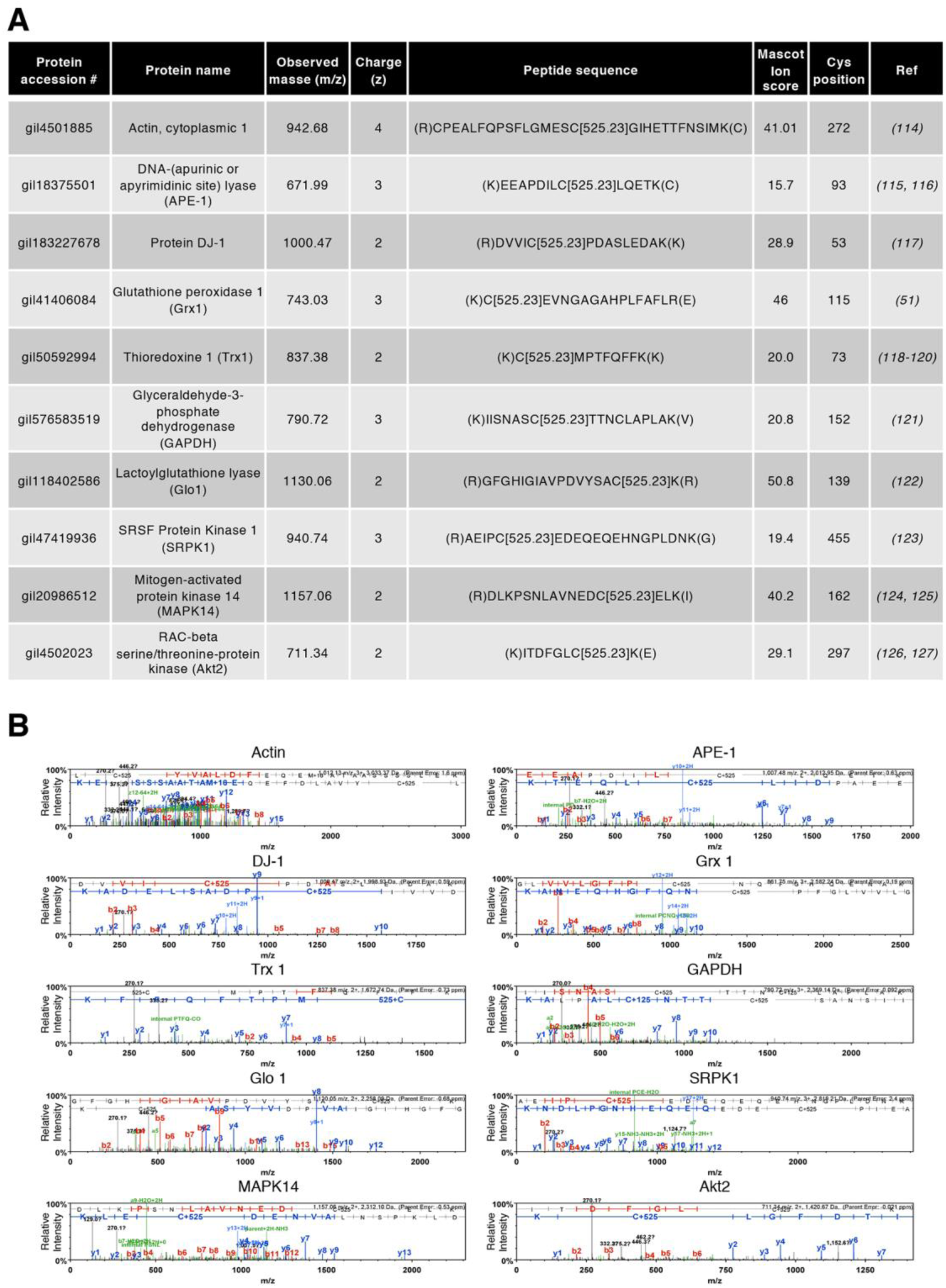
Validation of the InsPEx bioswitch method. **A**. Table showing Cys ox-PTMs sites that were previously documented in the literature and were also identified in this study. References: Actin-1 (*114*), APE-1 (*115, 116*), DJ-1 (*117*), Grx1 (*51*), Trx1 (*118-120*), GAPDH (*121*), Glo1 (*122*), SRPK1 (*123*), MAPK14 (*124, 125*), Akt2 (*126, 127*). **B**. Individual spectra of peptides described in (**A**). C+125 corresponds to a Cys alkylated with N-ethylmaleimide (NEM), thus, a Cys that was reduced in vivo. The fragment picks of both b and y ions are shown in the spectra in red and blue, respectively. All MS/MS spectra were visualized with scaffold, version 4.8.4.

**Figure S3.**
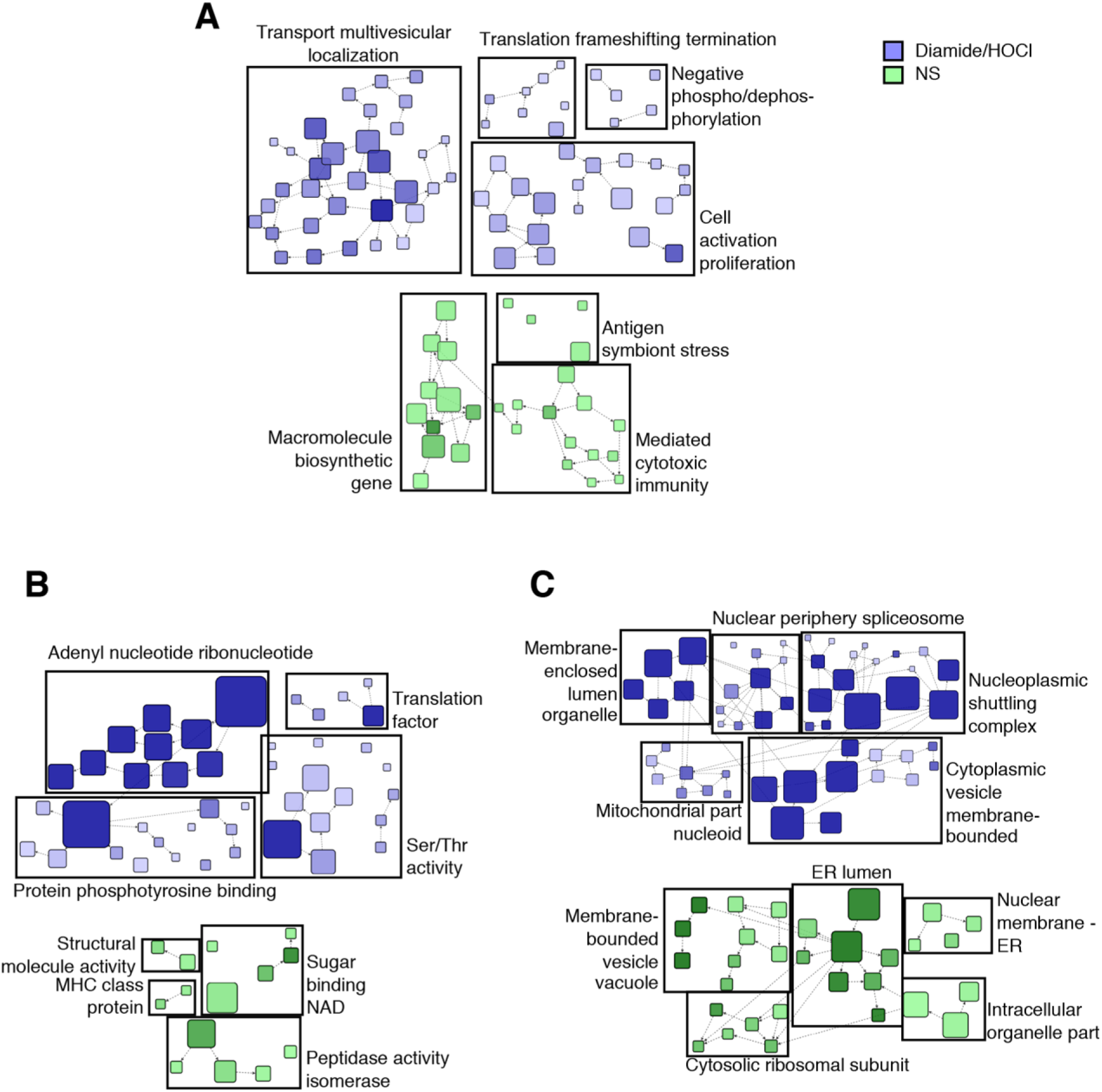
Community analysis of the GO terms enriched in Cys ox-PTMs. *Supplemental figure related to Fig. 2A-C*. Cluster visualization of the functional interaction networks generated by the fast greedy (GLay) algorithm. GLay was applied on networks in order to assemble the enriched biological processes (**A**), molecular functions (**B**) and cellular components (**C**). Terms enriched in non-stimulated (basal) and oxidant (diamide/HOCl)-treated conditions are shown in green and blue, respectively. The nodes of the network correspond to each enriched GO term and the direction of the edges (arrows) refers to their level of hierarchy. Arrows go from higher to lower hierarchy levels.

**Figure S4.**
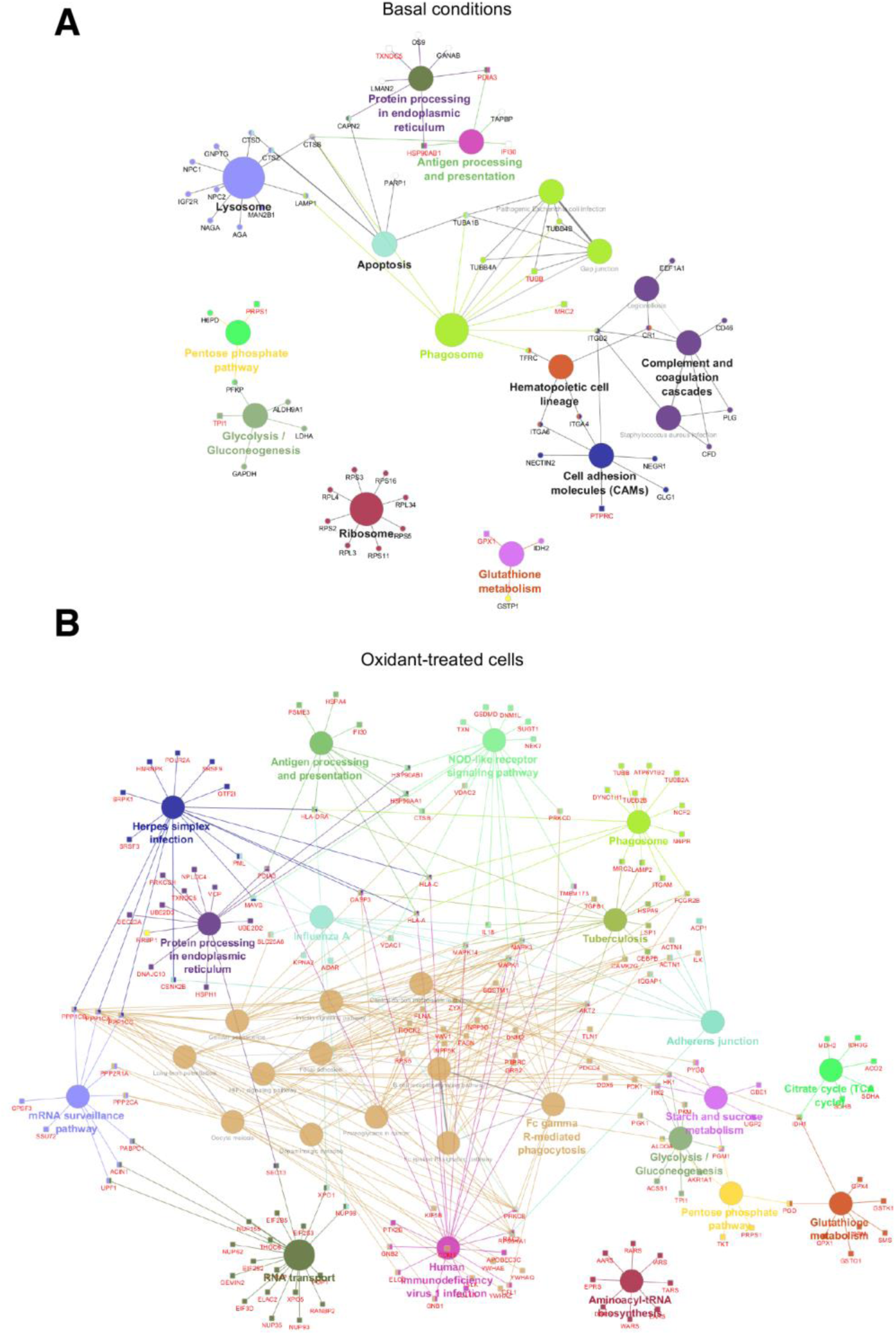
Complete map of all enriched KEGG pathways. *Supplemental figure related to Fig. 2D-F*. KEGG pathways enrichment analysis of identified reversible Cys ox-PTMs in non-stimulated (basal conditions) (**A**) and oxidant (diamide or HOCl)-stimulated (**B**) U937 cells. Oxidized proteins related with each pathway are indicated.

**Figure. S5.**
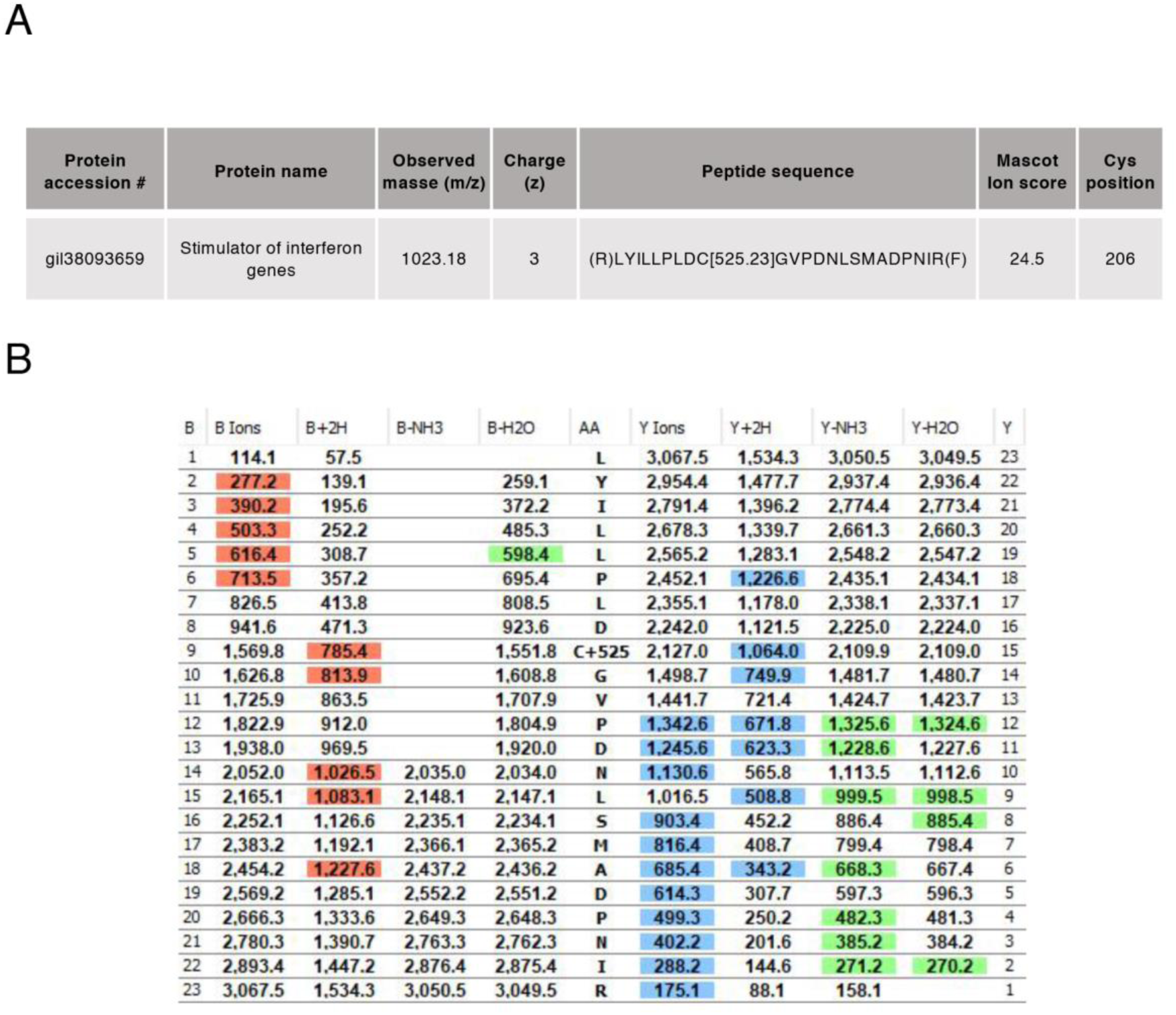
MS identification of STING Cys oxidation sites after 2’3’-cGAMP stimulation. *Supplemental figure related to Fig. 3F*. **A**. Table showing MS analysis of the identified STING peptide containing Mal-PEG_2_-Bio-labeled Cys^206^ (C[525.23]) after 2’3’-cGAMP stimulation. Peptides containing Cys^206^ were not detected in non-stimulated cells. **B**. Fragmentation table showing all b (red) and y (blue) ion values for the peptide containing the labeled Cys residue shown in (A). All MS/MS spectra were visualized with scaffold, version 4.8.4.

**Figure S6.**
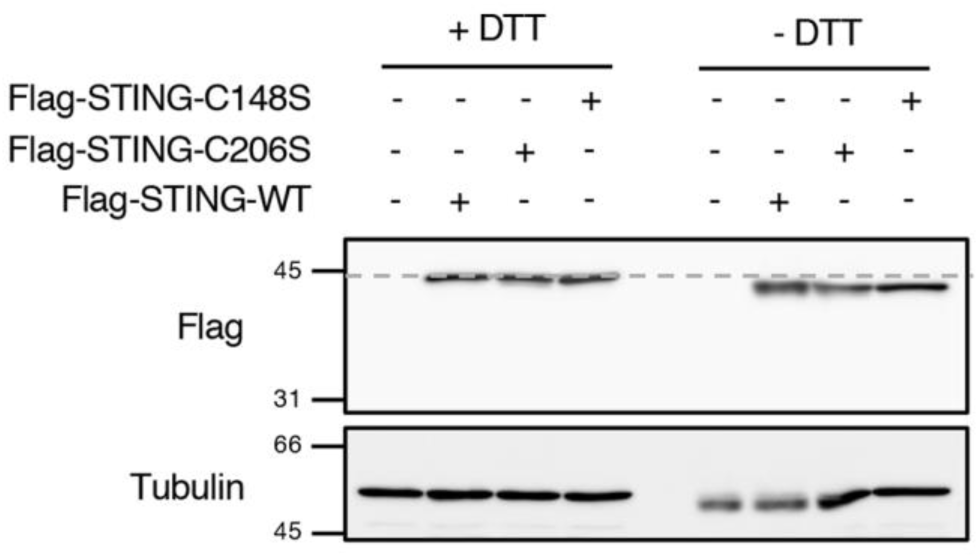
STING forms intramolecular disulfide bonds independently of Cys^148^ and Cys^206^. **A**. A549 cells were transfected with WT, C206S or C148S Flag-tagged STING encoding constructs. Whole Cell Extracts were resolved by reducing (with DTT) and non-reducing (without DTT) SDS-PAGE. Proteins were detected by immunoblot using anti-tubulin and anti-Flag antibodies. Molecular weight (kDa) markers are indicated on the left side of the SDS-PAGE gels.

**Figure S7.**
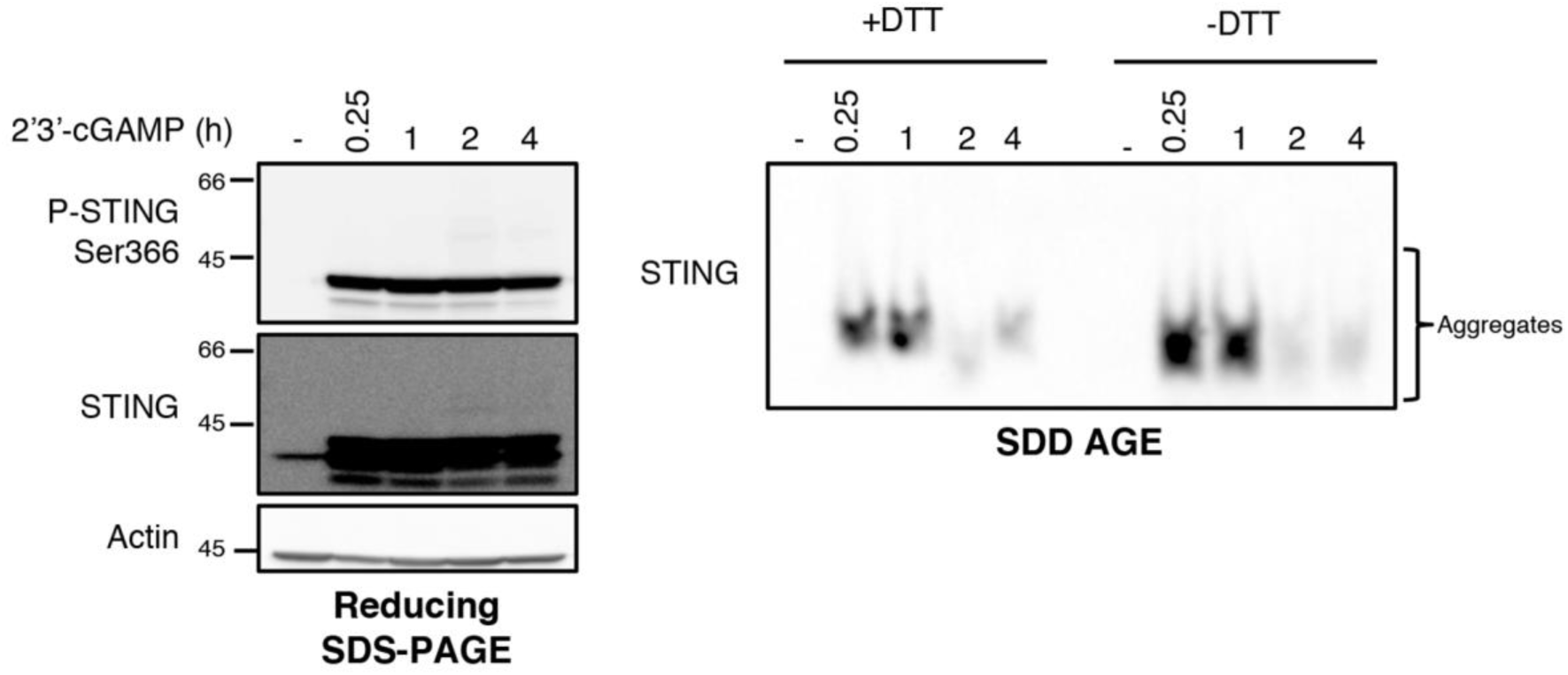
STING aggregates formed in response to 2’3’-cGAMP are sensitive to reducing agents. U937 cells were infused with 2’3’-cGAMP at 24µg/mL for the indicated times. Whole cell extracts were resolved by reducing (with DTT) and non-reducing (without DTT) SDD-AGE and by reducing SDS-PAGE. Proteins were detected by immunoblot using anti-actin, anti-STING and anti-phospho-STING-Ser^366^ antibodies. Molecular weight (kDa) markers are indicated on the left side of the SDS-PAGE gels.

**Figure S8.**
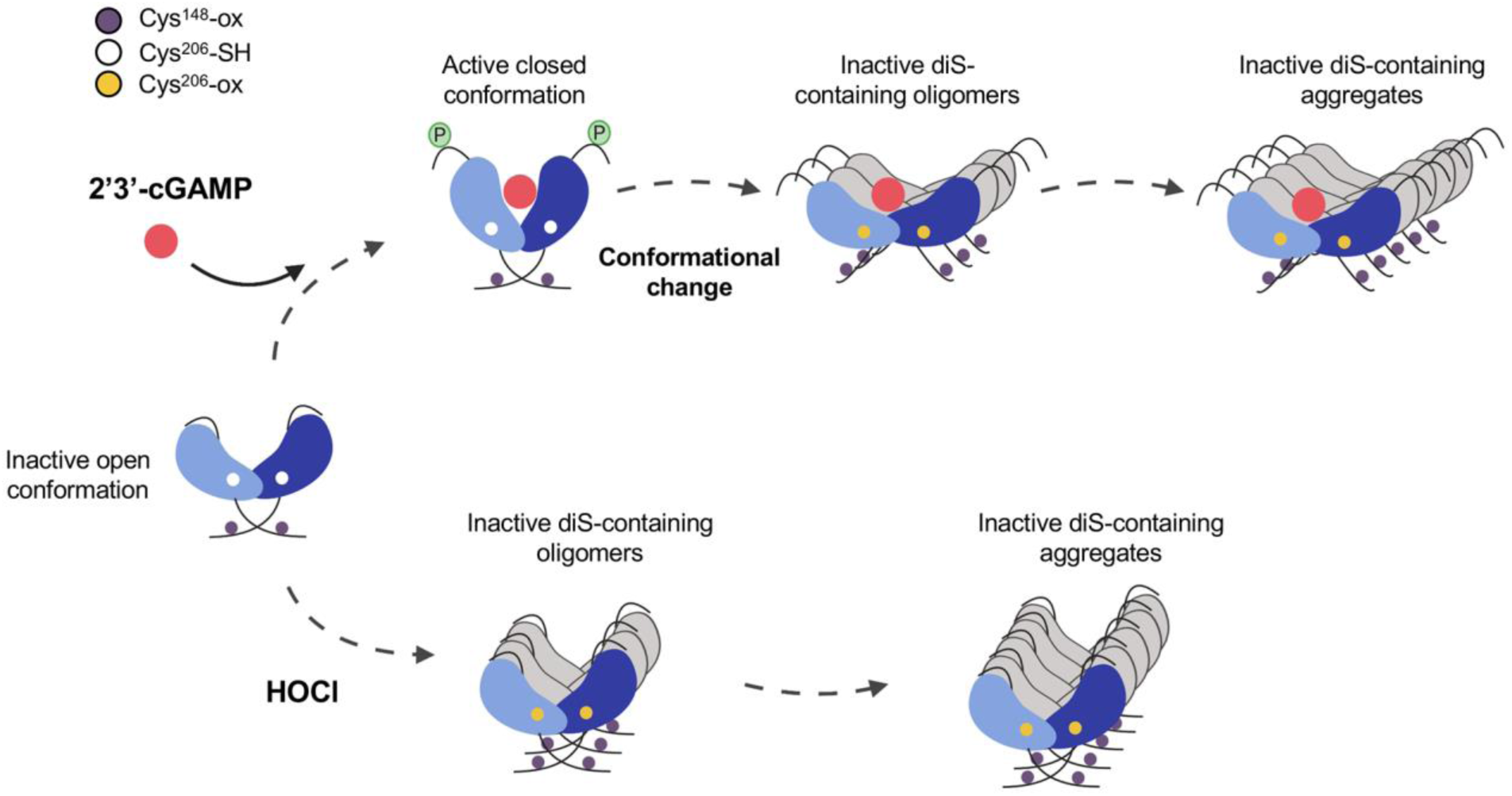
Model of STING regulation by Cys ox-PTMs. STING Cys^148^ is constitutively oxidized. Based on previous reports, this oxidation state is essential for STING to be activated by 2’3’-cGAMP (*61*). Binding of 2’3’-cGAMP triggers Ser^366^ phosphorylation and activation of STING. Furthermore, Cys^206^ undergoes oxidation, imposing a conformational change that leads to inhibition of Ser^366^ phosphorylation and provokes the formation of inactive diS-containing polymers to prevent STING hyperactivation. Oxidative stress induced by HOCl treatment without 2’3’-cGAMP stimulation, also triggers the formation of diS-containing polymers species.

